# SARS-CoV-2 nsp16 is regulated by host E3 ubiquitin ligases, UBR5 and MARCHF7

**DOI:** 10.1101/2024.08.30.610469

**Authors:** Li Tian, Zongzheng Zhao, Wenying Gao, Zirui Liu, Xiao Li, Wenyan Zhang, Zhaolong Li

**Author notes:** These authors contributed equally to this article (Li Tian, Zongzheng Zhao). Correspondence: Zhaolong Li, Wenyan Zhang, and Xiao Li.

## Abstract

Severe acute respiratory syndrome coronavirus 2 (SARS-CoV-2), the causative agent of coronavirus disease 2019 (COVID-19), remains a global public health threat with considerable economic consequences. The non-structural protein 16 (nsp16), in complex with nsp10, facilitates the final viral mRNA capping step through its 2′-O-methylase activity, helping the virus to evade host immunity and prevent mRNA degradation. However, nsp16 regulation by host factors remains poorly understood. While various E3 ubiquitin ligases interact with SARS-CoV-2 proteins, their roles in targeting nsp16 for degradation are unclear. In this study, we demonstrate that nsp16 undergoes ubiquitination and proteasomal degradation mediated by the host E3 ligases UBR5 and MARCHF7. UBR5 induces K48-linked ubiquitination, whereas MARCHF7 promotes K27-linked ubiquitination, independently suppressing SARS-CoV-2 replication in cell cultures and in mice. Notably, UBR5 and MARCHF7 also degrade nsp16 variants from different viral strains, exhibiting broad-spectrum antiviral activity. Our findings reveal novel antiviral mechanisms of the ubiquitin-proteasome system (UPS) and highlight their potential therapeutic targets against COVID-19.

## Introduction

Severe acute respiratory syndrome coronavirus 2 (SARS-CoV-2) remains a global public health threat, with over 777 million coronavirus disease 2019 (COVID-19) cases and 7.1 million deaths as of March 2025 (reported by World Health Organization). The virus encodes four structural proteins, 16 non-structural proteins (nsps) involved in replication and transcription, and several accessory proteins (ORFs) linked to immune evasion (V’kovski et al., 2021; Wang et al., 2020). Many viruses, including Ebola (Valle et al., 2021), dengue (Jung et al., 2018), and reovirus (Furuichi et al., 1975), encode 2’-O methyltransferases (2’-O-MTases) that mimic the host 5’ cap structure to evade innate immune recognition (Daffis et al., 2010; Züst et al., 2011). Similarly, SARS-CoV-2 nsp16, in complex with nsp10, functions as a 2′-O-MTase, modifying capped viral RNA from ‘cap-0’ to ‘cap-1’ (Benoni et al., 2021; Park et al., 2022). This modification protects the virus from host antiviral defences, such as MDA5 recognition and IFIT1 restriction (Bergant et al., 2022; Russ et al., 2022). Beyond immune evasion, nsp16 disrupts global host mRNA splicing, reducing cellular protein and mRNA levels (Banerjee et al., 2020). Nsp16 also enhances SARS-CoV-2 entry by upregulating TMPRSS2 expression (Han et al., 2023), underscoring its role as a virulence factor. Notably, nsp16-deficient strains exhibit low pathogenicity and elicit robust immune responses, suggesting their potential as live-attenuated vaccines (Ye et al., 2022). Given the critical functions of nsp16, inhibitors targeting nsp16 or the nsp16`-`nsp10 complex have been developed (Klima et al., 2022; Nguyen et al., 2022). Therefore, understanding host factors that regulate nsp16 is essential for identifying new therapeutic strategies.

The ubiquitin-proteasome system (UPS) is a key pathway for targeted protein degradation, often described as ‘the molecular kiss of death’ (Dongdem & Wezena, 2021; Park et al., 2020). Substrate ubiquitination involves a cascade of three enzymes: E1 activating enzyme, E2 conjugating enzyme, and E3 ubiquitin ligase (Zheng & Shabek, 2017). Some E3 ubiquitin ligases function as monomers, whereas others assemble into multi-subunit complexes. Regardless of their structure, all E3 ligases play a crucial role in substrate recognition, dictating UPS specificity. In certain complexes, an additional protein may function as a substrate receptor, mediating substrate-specific recognition (Iconomou & Saunders, 2016; Yang et al., 2021; Li et al., 2021; Wang et al., 2022). Over 600 E3 ligases encoded in the human genome regulate diverse cellular processes in response to various signals (Dongdem & Wezena, 2021; Garcia-Barcena et al., 2020). Dysregulation of their activity is linked to multiple diseases, making them attractive therapeutic targets (Humphreys et al., 2021). Based on conserved domains and ubiquitin (Ub) transfer mechanisms, E3 ligases are categorised into three classes: Really Interesting New Gene (RING), homologous to the E6AP carboxyl terminus (HECT), and RING-between-RING (RBR) ligases (Garcia-Barcena et al., 2020; Morreale & Walden, 2016). RING E3s transfer Ub directly from the E2-Ub complex to substrates (Deshaies & Joazeiro, 2009), whereas HECT-type E3s first attach Ub to their own catalytic cysteine before transferring it (Huibregtse et al., 1995). These distinct mechanisms highlight the complexity and versatility of E3 ligase function in cellular regulation (Walden & Rittinger, 2018; Garcia-Barcena et al., 2020).

To investigate the relationship between SARS-CoV-2 and host factors, we focused on UPS proteins that interact with nsp16. In this study, we demonstrate for the first time that nsp16 undergoes ubiquitination and proteasomal degradation. Notably, two E3 ligases from distinct families—RING-type MARCHF7 and HECT-type UBR5—independently target nsp16 for degradation, disrupting its function and exhibiting strong antiviral activity against SARS-CoV-2. Our findings reveal novel therapeutic targets for combating SARS-CoV-2 and treating COVID-19.

## Results

### SARS-CoV-2 nsp16 is degraded via the ubiquitin-proteasome pathway

Ubiquitination plays a crucial role in SARS-CoV-2 infection and pathogenesis (Gao et al., 2022; Guo et al., 2021; Li et al., 2023; Zhang et al., 2021; Zhang et al., 2024; Zhang et al., 2023). To determine whether SARS-CoV-2 non-structural proteins (nsps) are UPS-regulated, we examined the effect of the proteasome inhibitor, MG132, on nsps expression. MG132 treatment significantly increased the nsp8, nsp11, and nsp16 abundance in HEK293T cells (Fig. 1A), suggesting UPS-mediated degradation. Nsp8 undergoes TRIM22-mediated ubiquitination and proteasomal degradation (Fan et al., 2024). In this study, we focused on nsp16 for further investigation. To confirm UPS-mediated nsp16 degradation, we tested additional proteasome inhibitors, Bortezomib and Carfilzomib, both of which enhanced nsp16 stability. In contrast, lysosomal inhibitors (Bafilomycin A1 and NH_4_Cl) and the autophagy-lysosomal inhibitor, Vinblastine, had no such effect (Fig. 1B). To further assess the role of the UPS in regulating nsp16 stability, we treated nsp16-expressing cells with cycloheximide, a protein synthesis inhibitor, and measured the half-life of nsp16. MG132 treatment significantly prolonged nsp16 stability, extending its half-life from 2 h to 15 h (Fig. 1C and 1D).

**Figure 1.**
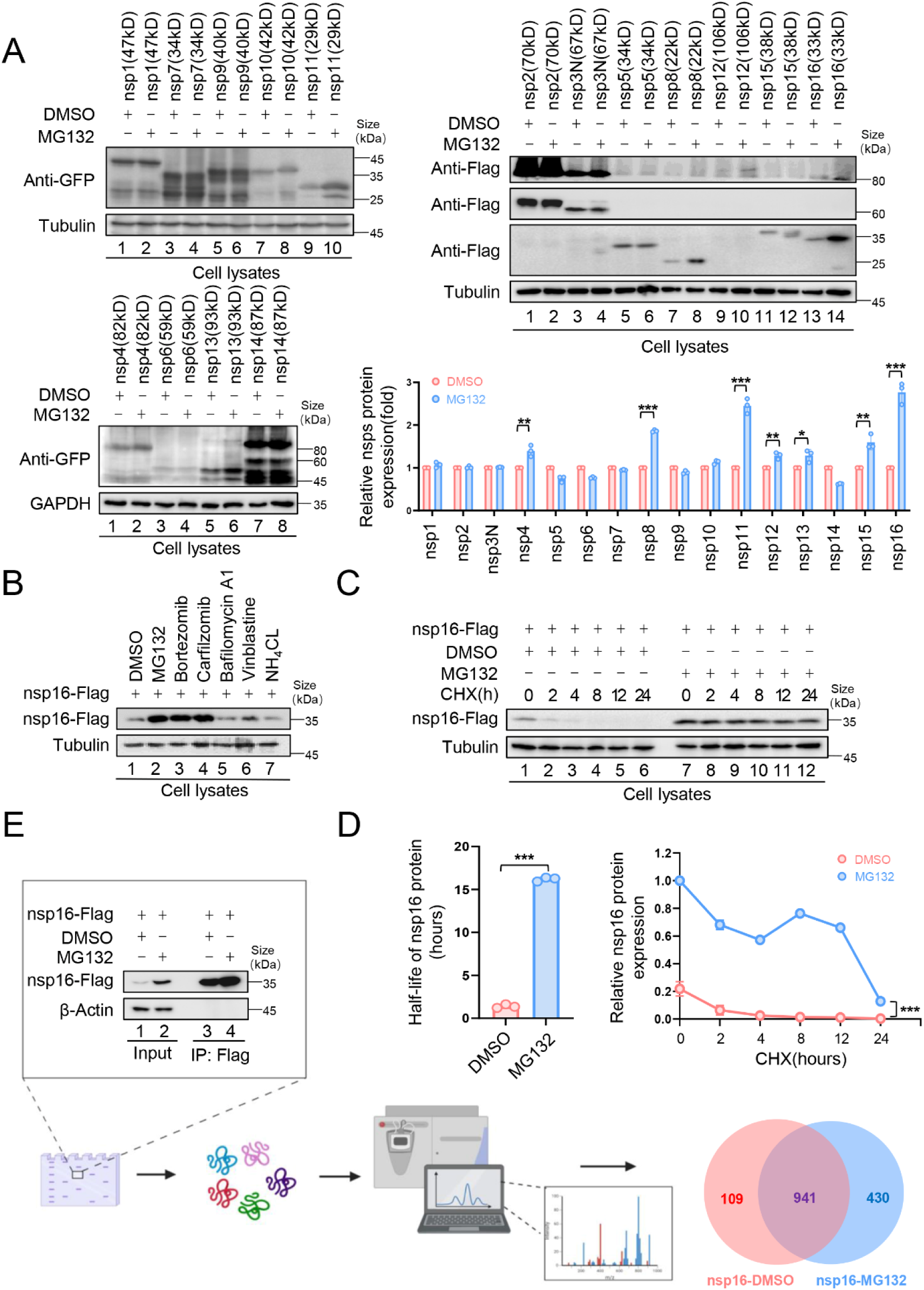
The non-structural protein nsp16 of SARS-CoV-2 was identified that can be degraded through the proteasome pathway. A. The non-structural proteins nsp8, nsp11 and nsp16 could be restored by the proteasome inhibitor MG132. HEK293T cells in 12-well plates were transfected with the plasmids of 16 nonstructural proteins (nsp1-16) encoded by SARS-CoV-2. Thirty-six hours later, the cells were treated with MG132 (10 µM) or DMSO for 12 h before collection. The protein level was detected by Immunoblotting (IB). Quantification of nsp protein levels relative to the control protein is shown. Data are representative of three independent experiments and shown as average ±SD (n = 3). Significance was determined by a two-tailed t-test: *P < 0.05; **P < 0.01; ***P < 0.001. B. Proteasomal inhibitors but no other inhibitors stabilized nsp16 protein. HEK293T cells transfected with the nsp16-Flag expression vector were treated with dimethyl sulfoxide (DMSO), MG132 (10 µM), Bortezomib (10 µM), Carfilzomib (10 µM), Bafilomycin A1 (5 µM), Vinblastine (2.5 µM), or NH_4_CL (2.5 µM) for 12 h prior to harvest. The cell lysates were analyzed by anti-Flag antibody. (C-D). The half-life of nsp16 was prolonged by the proteasome inhibitor MG132. C. HEK293T cells were transfected with the nsp16-Flag-expressing plasmids. 12 hours later, the cells were treated with DMSO or MG132 (10 µM) for 12 h, then 50 µg/mL cycloheximide (CHX) was added. Cells were harvested at the indicated times to detect the level of viral protein by anti-Flag antibody. D. Quantification of nsp16 protein levels relative to tubulin at different time points is shown. The half-life of the nsp16 protein was determined based on protein quantification using Image J, combined with the protein half-life formula for calculation. Results are shown as mean ± SD (n = 3 independent experiments). ***, P < 0.001 by by a two-tailed t-test. E. Samples were prepared for mass spectrometry, and nsp16 interacting proteins were obtained by immunoprecipitation (IP) (created with BioRender.com and the agreement no. is LC27PCBEOF). The plasmids were transfected into HEK293T cells for 48 h. Treat cells with or without MG132 (10 µM) for 12 h prior to harvest. The whole-cell lysates were incubated with protein G agarose beads conjugated with anti-Flag antibodies and used for IB with anti-Flag antibodies to detect the nsp16 protein. Samples enriched for proteins were analyzed by mass spectrometry. Figure 1-source data 1. PDF file containing original western blots for Figure 1A, B, C, and E, indicating the relevant bands and treatments. Figure 1-source data 2. Original files for western blot analysis displayed in Figure 1A, B, C, and E. Figure 1-source data 3. Numerical data obtained during experiments represented in Figure 1.

To identify nsp16-interacting proteins, we performed co-immunoprecipitation (Co-IP) followed by mass spectrometry (MS) analysis, comparing nsp16-expressing cells with and without MG132 treatment (Fig. 1E). Kyoto Encyclopedia of Genes and Genomes (KEGG) pathway and Gene Ontology (GO) analyses (Figure 1—figure supplement 1A) revealed that inhibiting nsp16 degradation increased its interacting proteins, including 43 associated with viral processes, 22 involved in mRNA stability regulation, and five linked to cellular antiviral defence. Additionally, interactions related to mRNA splicing and RNA methylation were identified (Banerjee et al., 2020), consistent with the function of nsp16 as a 2′-O-MTase. Furthermore, SARS-CoV-2 exploits nsp16 to disrupt host mRNA splicing, which promotes infection (Park et al., 2022; Russ et al., 2022; Züst et al., 2011), further validating our MS data.

As expected, several nsp16-binding proteins were associated with ubiquitination and degradation pathways (Figure 1—figure supplement 1B). We identified four deubiquitinases (DUBs), the E2 ligase UBE2D3, and 14 proteasomal enzymes. Among them, six E3 ligases—UBR5, MARCHF7, HECTD1, TRIM32, MYCBP2, and TRIM21—were selected for further investigation.

### E3 ligases, UBR5 and MARCHF7, independently mediate nsp16 degradation

To identify the E3 ligases responsible for nsp16 degradation, we designed small interfering RNAs (siRNAs) targeting six candidate E3 ligases and assessed their effects on nsp16 abundance. *UBR5* and *MARCHF7* knockdown significantly stabilised nsp16 protein levels (Fig. 2A). To further investigate their roles in nsp16 degradation, we generated stable cell lines with *UBR5* or *MARCHF7* knockdowns and confirmed silencing efficiency (Fig. 2B). To determine whether UBR5 and MARCHF7 function cooperatively, we performed dual knockdowns. Silencing *UBR5* in *MARCHF7*-knockdown cells, and vice versa, further enhanced nsp16 stability, indicating that these ligases act independently (Fig. 2C). Overexpression studies further supported their independent roles—MARCHF7 overexpression induced nsp16 degradation even in *MARCHF7*-knockdown and *MARCHF7/UBR5* double-knockdown cells. Similarly, UBR5 overexpression promoted nsp16 degradation despite *MARCHF7* knockdown. Silencing efficiencies were validated across all experiments (Figure 2—figure supplement 1A and 1B). Notably, wild-type MARCHF7 and UBR5, but not their mutants lacking functional RING (1-542 aa) or HECT domains, respectively, effectively degraded nsp16 in knockdown cells (Fig. 2D and 2E). These findings confirm that UBR5 and MARCHF7 independently mediate nsp16 ubiquitination via their HECT and RING domains, targeting it for proteasomal degradation.

**Figure 2.**
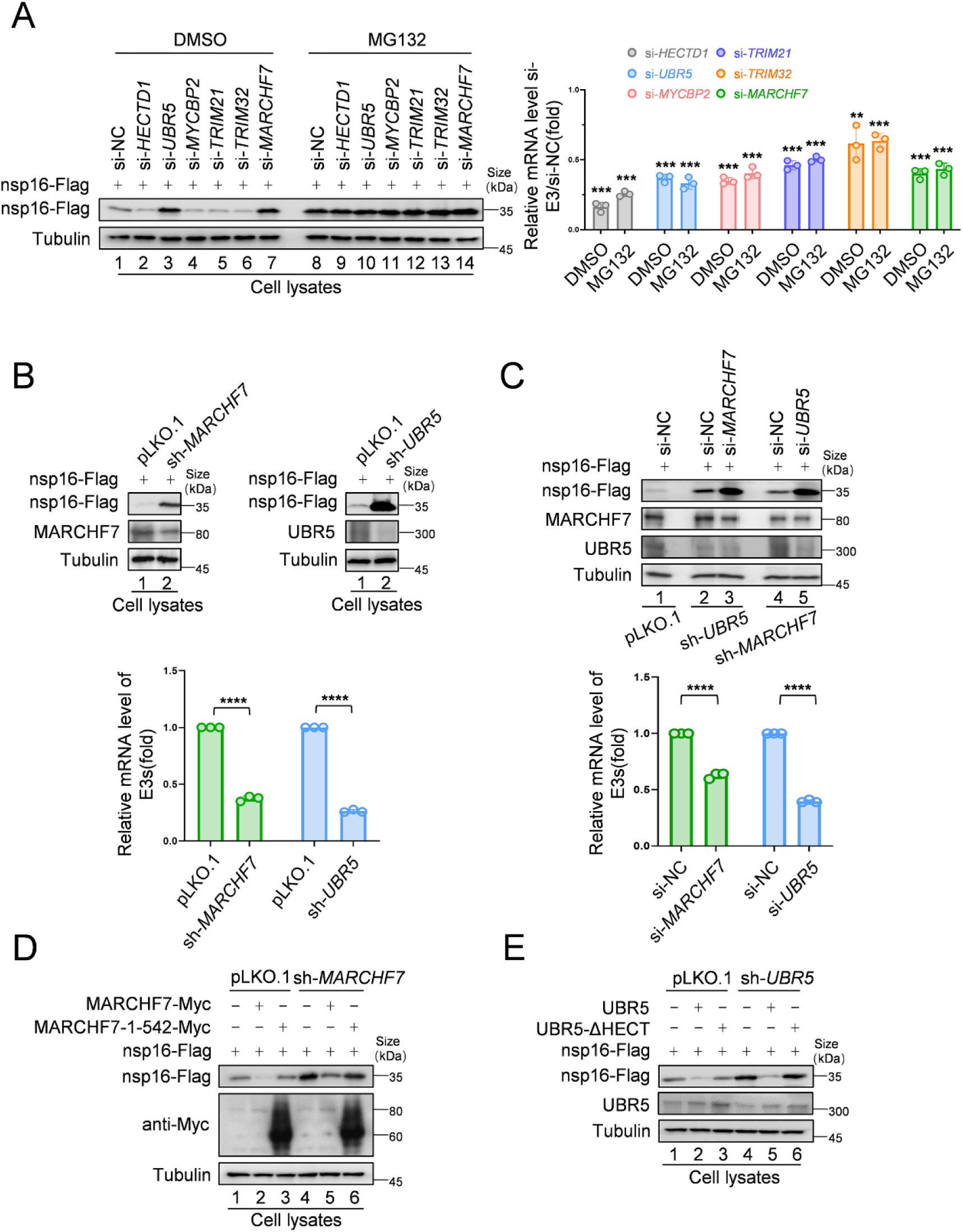
MARCHF7 and UBR5 were identified as E3 ubiquitin ligases involves in nsp16 protein degradation. A. Knockdown of *MARCHF7* or *UBR5* resulted in nsp16 restoration. HEK293T cells were transfected with siRNA of E3 ligase candidates for 24 h, followed by co-incubation with the nsp16-Flag-expressing plasmids for 48 h, treated with MG132 (10 µM) for 16h before harvesting, lysed, and subjected to IB assay using anti-Flag antibody. RT-qPCR was conducted to determine the mRNA expression levels of E3 ligase candidates. The si-RNA targeting regions for the candidate E3 ubiquitin ligase proteins and the targeted regions for RT-qPCR are shown in Figure 2—figure supplement 1C. Data are representative of three independent experiments and shown as average ± SD (n = 3). Significance was determined by a two-tailed t-test: ***P < 0.001. B. RNA levels of UBR5 or MARCHF7 from HEK293T cells infected with lentivirus containing control or shRNA targeting *UBR5* or *MARCHF7* for 48 h and screened with antibiotics for 48 h. Knockdown cell lines were transfected with plasmids expressing nsp16-Flag, collected at the indicated times, and the protein levels of nsp16, MARCHF7, and UBR5 were detected by IB.C. MARCHF7 and UBR5 acted separately and did not depend on each other. HEK293T cells stably expressing *UBR5* shRNA or *MARCHF7* shRNA were transfected with siRNA of *MARCHF7* or *UBR5* for 24 h, respectively, followed by co-incubation with the nsp16-Flag-expressing plasmids for 48 h. The protein levels and the RNA levels of nsp16, UBR5 and MARCHF7 were measured by IB and RT-qPCR, respectively. (D-E). In HEK293T cells stably expressing *UBR5* shRNA or *MARCHF7* shRNA, nsp16 was degraded by overexpressed UBR5 or MARCHF7, respectively, whereas the mutant failed to degrade nsp16. The cell lysates were analyzed by anti-Flag antibody. Figure 2-source data 1. PDF file containing original western blots for Figure 2A-E, indicating the relevant bands and treatments. Figure 2-source data 2. Original files for western blot analysis displayed in Figure 2A-E. Figure 2-source data 3. Numerical data obtained during experiments represented in Figure 2.

### UBR5 and MARCHF7 mediate distinct Ub linkages on nsp16

We first confirmed nsp16 ubiquitination both with and without exogenous ubiquitin (Ub) expression (Fig. 3A-B). To investigate the role of E3 ligases in this process, we examined the impact of *UBR5* and *MARCHF7* knockdown on nsp16 ubiquitination. Silencing either ligase reduced nsp16 ubiquitination compared to the negative control, indicating their involvement in this modification (Fig. 3C). Ub contains seven lysine residues (K6, K11, K27, K29, K33, K48, and K63), which form polyubiquitin chains with distinct functions that determine the fate of the attached protein (Grice & Nathan, 2016). To identify the specific polyubiquitin chain types formed on nsp16 by UBR5 and MARCHF7, we used a series of Ub mutants, each retaining only a single lysine residue. Except for K33, all single-lysine Ub mutants supported nsp16 ubiquitination to varying degrees, suggesting a complex ubiquitin architecture potentially regulated by multiple E3 ligases or E2-E3 pairs (Fig. 3D). Further analysis of lysine-specific linkages revealed that MARCHF7 primarily mediates K27-linked ubiquitination of nsp16, whereas UBR5 facilitates K48-linked ubiquitination (Fig. 3E and 3F). These findings establish that UBR5 and MARCHF7 independently regulate nsp16 degradation through distinct ubiquitin linkages.

**Figure 3.**
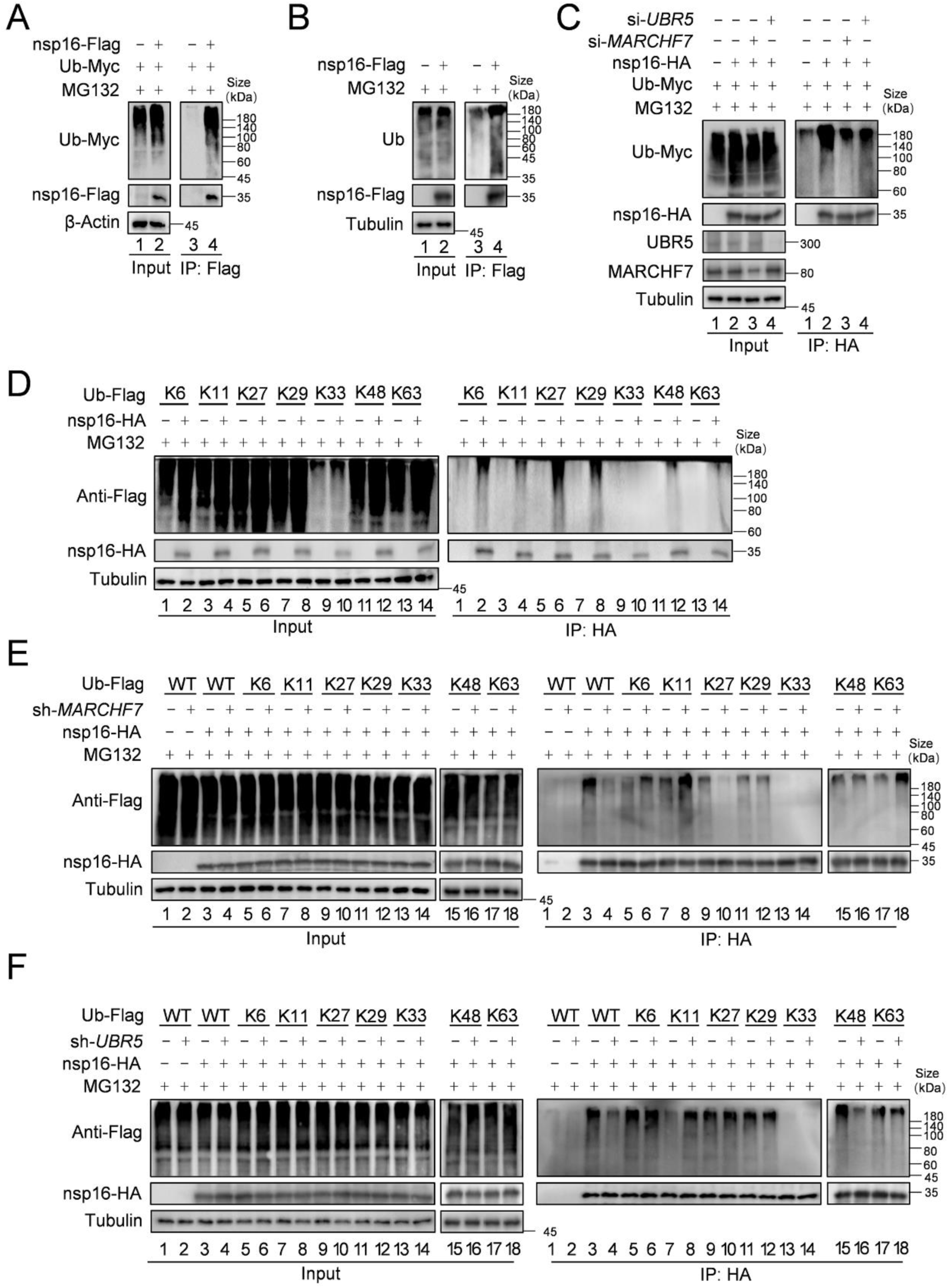
MARCHF7 or UBR5 catalyze the formation of K-27 type or K-48 type ubiquitin chains of nsp16 respectively. A. Nsp16 can be ubiquitinated. HEK293T cells co-transfected with ubiquitin-Myc and nsp16-Flag or transfected with nsp16-Flag alone. The cells were treated with MG132 for 12 h before collection. The whole-cell lysates were incubated with anti-Flag beads and used for IB with anti-Myc or anti-Flag antibodies to detect the polyubiquitination chain of nsp16. B. Assess the endogenous ubiquitination level of nsp16 protein. Cells were transfected with nsp16-Flag or an empty vector, and collected 48 hours later. Prior to harvesting, cells were treated with MG132 for 16 hours. Co-IP experiments were then performed to analyze the endogenous ubiquitination level of nsp16. C. The level of ubiquitination of nsp16 decreased with decreasing the protein levels of MARCHF7 or UBR5. E3 was knocked down by transfection with siRNA targeting *UBR5* or *MARCHF7*, and 24 h later ubiquitin-Myc and nsp16-HA were co-transfected or nsp16-HA alone. Cells were treated with MG132 for 16 h before collection. Whole cell lysates were incubated with anti-HA beads, and polyubiquitinated chains of nsp16 were detected by IB with anti-Myc or anti-HA antibodies. D. Nsp16 can be modified by a variety of ubiquitin chains. HEK293T cells were transfected with either nsp16-HA alone or together with plasmids encoding various mutants of ubiquitin (K6 only, K11 only, K27 only, K29 only, K33 only, K48 only, K63 only). Thirty-six hours later, cells were treated with MG132 for 12 h. Cell lysates were then subjected to immunoprecipitation, followed by IB to analysis. (E-F) MARCHF7 or UBR5 causes nsp16 to be modified by the K27 type or K48 type ubiquitin chain. 293T cell lines with or without *MARCHF7* or *UBR5* knockdown were co-transfected with plasmids encoding ubiquitin-WT or various mutants of ubiquitin (K6 only, K11 only, K27 only, K29 only, K33 only, K48 only, K63 only). The other experimental methods were the same as C. Figure 3-source data 1. PDF file containing original western blots for Figure 3A-F, indicating the relevant bands and treatments. Figure 3-source data 2. Original files for western blot analysis displayed in Figure 3A-F.

### UBR5 and MARCHF7 directly interact and colocalise with nsp16 in the endoplasmic reticulum (ER)

MS analysis suggested interactions between nsp16 and both UBR5 and MARCHF7. Co-IP experiments confirmed that Myc-tagged MARCHF7 and endogenous UBR5 bind to nsp16 (Figure 4—figure supplement 1A and 1B). To assess whether these interactions are direct, we performed fluorescence resonance energy transfer (FRET) assays. When nsp16-YFP was bleached, the fluorescence intensity of CFP-UBR5 and CFP-MARCHF7 increased, confirming direct interactions (Figure 4—figure supplement 1C). ImageJ software was used to quantify relative fluorescence intensities (Figure 4—figure supplement 1D). To determine whether UBR5 and MARCHF7 depend on each other for nsp16 binding, we examined the effect of *MARCHF7* knockdown on UBR5-nsp16 interactions, and vice versa, using Co-IP. The interactions remained unchanged, indicating that UBR5 and MARCHF7 independently bind to nsp16 (Fig. 4A and 4B). Nsp16 localises to both the nucleus and cytoplasm (Zhang et al., 2020). UBR5 (containing two nuclear localization signals) and MARCHF7 are also present in both compartments (Muñoz-Escobar et al., 2015; Shearer et al., 2018) (Nathan et al., 2008). Immunofluorescence staining revealed that UBR5 and MARCHF7 colocalised with nsp16 predominantly in the cytoplasm, with some nuclear colocalization observed in HeLa cells (Figure 4—figure supplement 1E). Similar results were obtained in nsp16-transfected HEK293T cells using antibodies against endogenous UBR5 and MARCHF7 (Figure 4—figure supplement 1F).

**Figure 4.**
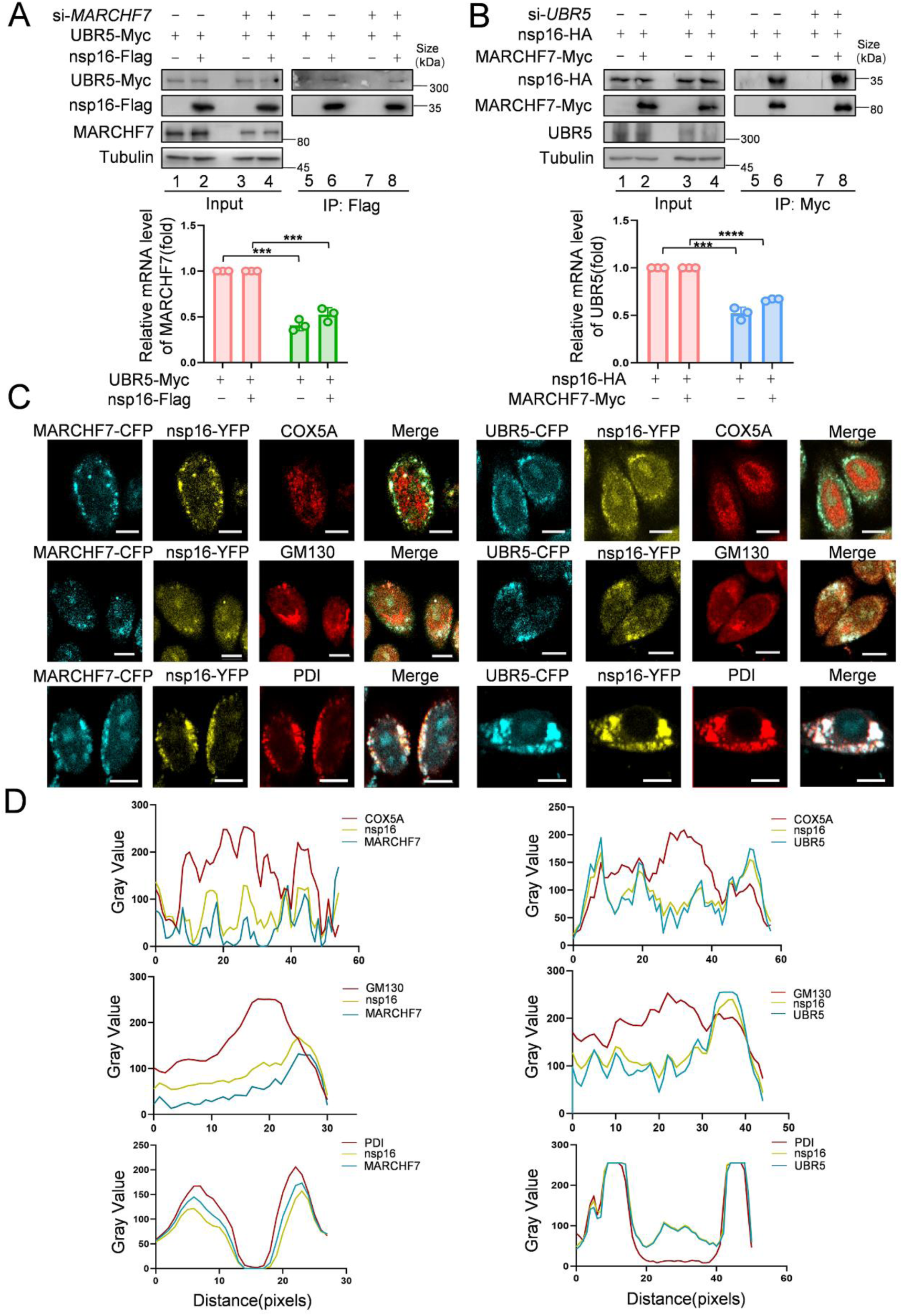
MARCHF7 and UBR5 directly interact with nsp16 respectively. (A-B). The binding of MARCHF7 or UBR5 to nsp16 was not mutually dependent. The binding of nsp16 to UBR5 or MARCHF7 was identified by co-immunoprecipitation in HEK293T cells transfected si*MARCHF7* or si*UBR5*, respectively. The immunoprecipitates and input were analyzed by IB. The knockdown efficiency was detected by RT-qPCR and IB. (C-D). MARCHF7 or UBR5 co-localized with nsp16 in the endoplasmic reticulum. Hela cells were co-transfected with YFP-nsp16(yellow) and CFP-UBR5(cyan) or CFP-MARCHF7(cyan). The organelles were labeled with antibodies against marker proteins of endoplasmic reticulum, Golgi apparatus and mitochondria respectively(red). The cells were analyzed by confocal microscopy (C). Scale bars, 20 um. The ratio of colocalization was quantified by measuring the fluorescence intensities using Image J (D). Figure 4-source data 1. PDF file containing original western blots for Figure 4A and B, indicating the relevant bands and treatments. Figure 4-source data 2. Original files for western blot analysis displayed in Figure 4A and B. Figure 4-source data 3. Numerical data obtained during experiments represented in Figure 4.

To pinpoint the specific cellular compartment where these interactions occur, we co-transfected UBR5-CFP or MARCHF7-CFP with nsp16-YFP and performed immunostaining using organelle-specific markers: COX5A (mitochondria), PDI (ER), and GM130 (Golgi apparatus). Notably, UBR5 and MARCHF7 both interacted with nsp16 and colocalised with PDI, but not with COX5A or GM130 (Fig. 4C and 4D), indicating that both E3 ligases interact with nsp16 primarily in the ER.

### Functional domains of UBR5 and MARCHF7 are required for nsp16 interaction and ubiquitination

UBR5 is a four-domain E3 ligase containing two nuclear localization signals. Its domains include UBA, UBR, PABC, and HECT (Muñoz-Escobar et al., 2015) (Figure 4—figure supplement 2A). Notably, the HECT domain is critical for its E3 ligase activity, as UBR5 must first conjugate to Ub before transferring it to substrates (Kim et al., 2021). To determine which UBR5 domain mediates nsp16 ubiquitination, we used UBR5 mutants with individual domain inactivation. Only the HECT domain mutant failed to degrade nsp16 (Figure 4—figure supplement 2B), consistent with previous studies (Zhou et al., 2022). This mutant also lost the ability to ubiquitinate nsp16, confirming the essential role of the HECT domain in this process (Figure 4—figure supplement 2C).

To identify the MARCHF7 region responsible for nsp16 degradation, we generated a series of truncation mutants as described previously (Figure 4—figure supplement 2D) (Zhao et al., 2018) (Nathan et al., 2008). All mutants lost the ability to degrade nsp16 (Figure 4—figure supplement 2E). Further analysis revealed that only the mutant containing the intact N-terminal region (aa 1–542) strongly bound to nsp16, while those retaining the active RING domain region (aa 543–616) did not. Consistently, only wild-type MARCHF7 promoted K27-linked ubiquitination of nsp16 (Figure 4—figure supplement 2F). These results indicate that the N-terminal region of MARCHF7 is essential for nsp16 binding, whereas its RING domain is required for K27-linked ubiquitination activity.

### UBR5 and MARCHF7 suppress SARS-CoV-2 replication by targeting nsp16 for degradation

To assess the impact of nsp16 degradation on viral replication, we used a biosafety level 2 system to generate SARS-CoV-2 transmissible virus-like particles capable of infecting Caco2-N^int^ cells (Ju et al., 2021). Knockdown of either *UBR5* or *MARCHF7* significantly increased SARS-CoV-2 replication, with simultaneous knockdown further enhancing viral replication (Figure 5—figure supplement 1A and 1B).

To investigate nsp16 modifications following infection, we infected HEK293T-ACE2 cells with the IME-BJ01 strain for 48 h. Co-IP using an anti-nsp16 antibody confirmed its interaction with endogenous MARCHF7 and UBR5, along with ubiquitination (Fig. 5A). In a biosafety level 3 facility, we further validated the antiviral roles of UBR5 and MARCHF7 using the IME-BJ01 (MOI 0.01) and Omicron strains (MOI 0.001). Knockdown of either E3 ligase in Caco2 cells significantly increased intracellular and secreted viral mRNA levels (M and E genes) for both strains. Viral titres in the supernatants also increased, with *UBR5* knockdown causing over a 1-log rise for the IME-BJ01 strain and even greater effects for Omicron (Fig. 5C–E and 5F–H for the IME-BJ01 and Omicron strains, respectively). Immunoblotting confirmed increased intracellular and secreted N protein levels upon *UBR5* or *MARCHF7* knockdown (Fig. 5I), with knockdown efficiencies verified (Fig. 5B). Due to low transfection efficiency in Caco2 cells, we overexpressed UBR5-Myc or MARCH7-Myc in HEK293T-ACE2 cells. Overexpression significantly reduced viral mRNA (M and E genes) and N protein levels in both strains, accompanied by a 0.5-log decrease in viral titres. However, co-transfection with increasing nsp16 levels counteracted these inhibitory effects (Fig. 6A–H and Figure 6—figure supplement 1A–H). To confirm the role of UBR5 and MARCHF7 enzymatic activity, we overexpressed catalytically inactive mutants (UBR5-Δ HECT or MARCHF7-aa 1-542) alongside nsp16. These mutants failed to suppress viral replication. Additionally, gradual increases in nsp16 expression did not further enhance viral replication, with only a slight increase in M mRNA, while E and N protein levels remained unchanged (Figure 6—figure supplement 2A–H). These findings demonstrate that UBR5 and MARCHF7 exert antiviral effects by ubiquitinating and degrading nsp16, thereby significantly inhibiting SARS-CoV-2 replication.

**Figure 5.**
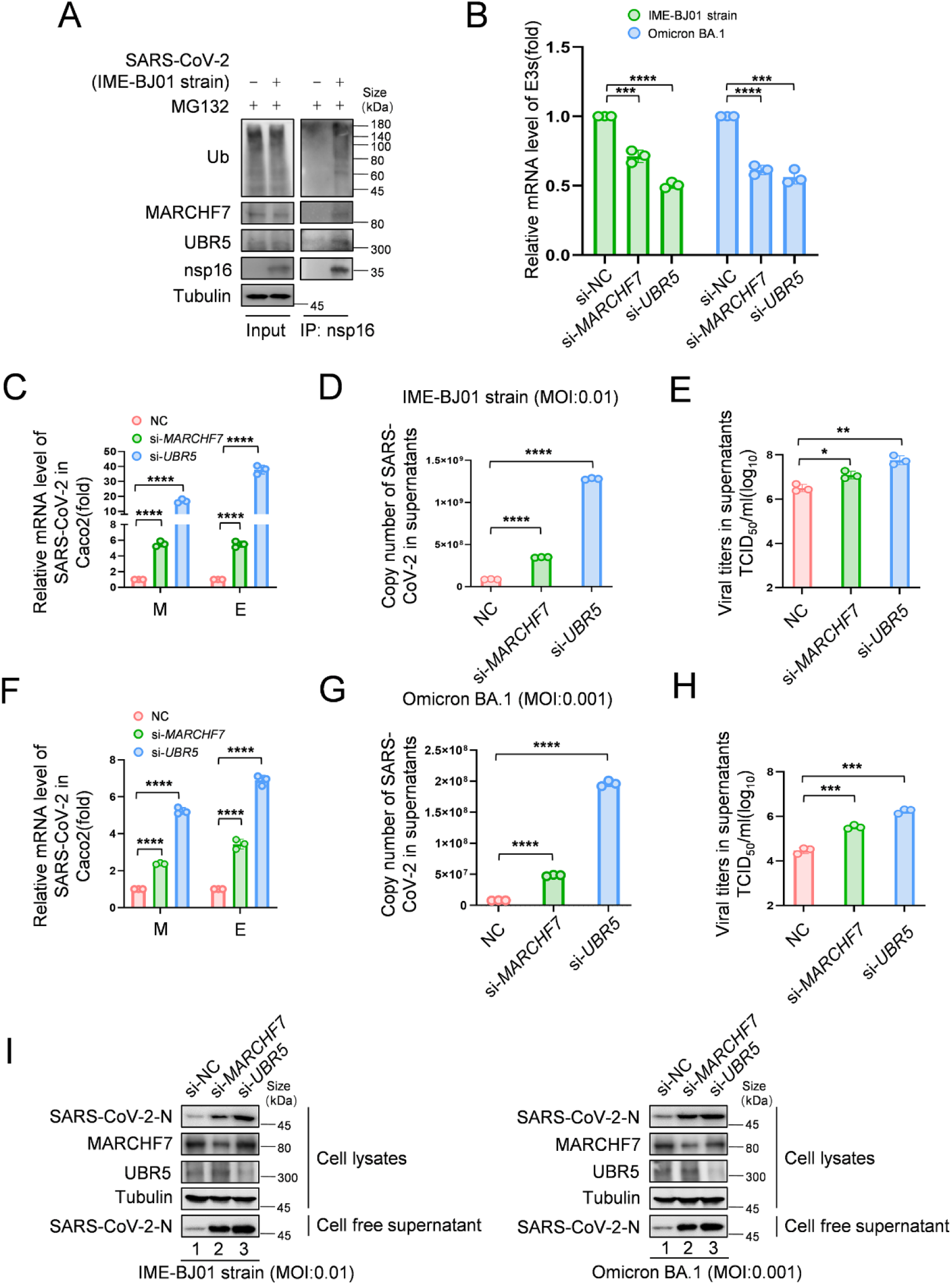
Knockdown of *MARCHF7* or *UBR5* promotes viral replication. (I). The virus-encoded nsp16 protein interacts with endogenous MARCHF7 and UBR5 and undergoes ubiquitination modification. In 293T-ACE2 cells, with or without IME-BJ01 strain infection (MOI: 0.01), the medium was changed 2 hours post-infection, and cells were harvested 48 hours later, with MG132 treatment added 16 hours before harvesting. nsp16 protein was enriched using Protein-G beads coupled with the nsp16 antibody, and interactions and ubiquitination were analyzed by immunoblotting (IB) with endogenous antibodies against MARCHF7, UBR5, and ubiquitination. (B-I). *MARCHF7* and *UBR5* were knocked down by siRNA in Caco2 cells. 24 h after transfection, the cells were infected with IME-BJ01 strain (MOI:0.01) (C-E) or Omicron BA.1 strain (MOI: 0.001) (F-H), respectively. 2 h post infection, the supernatant was discarded, and the cells were cultured in DMEM containing 3% fetal bovine serum for 48 h. The mRNA levels of SARS-CoV-2 M and E genes in the cells (C, F) and E genes in supernatant (D, G) were detected by RT-qPCR and the viral titers in supernatant (E, H) were measured. The N protein levels of IME-BJ01 or Omicron viruses were detected by IB (i). Knock-down efficiencies of *MARCHF7* and *UBR5* were detected by RT-qPCR or IB (B, I). Data are representative of three independent experiments and shown as average ±SD (n = 3). Significance was determined by one-way ANOVA, followed by a Tukey multiple comparisons posttest: *P < 0.05; **P < 0.01; ***P < 0.001. Figure 5-source data 1. PDF file containing original western blots for Figure 5A and I, indicating the relevant bands and treatments. Figure 5-source data 2. Original files for western blot analysis displayed in Figure 5A and I. Figure 5-source data 3. Numerical data obtained during experiments represented in Figure 5.

**Figure 6.**
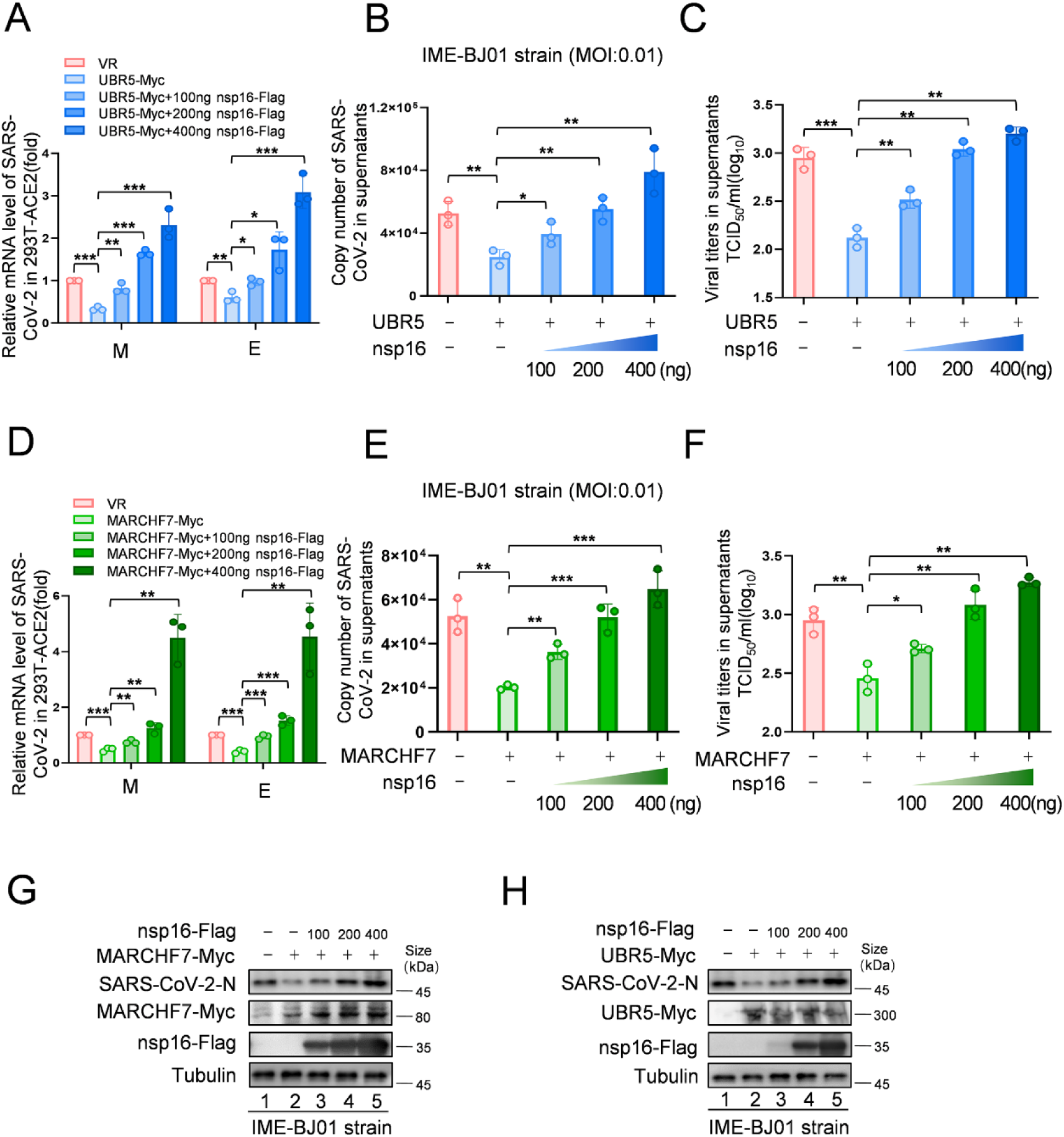
Increased levels of nsp16 rescued viral inhibition by UBR5 or MARCHF7’. (A-H) UBR5 or MARCHF7 was transfected in 293T cells stably overexpressed with ACE2, and the increased doses of nsp16-Flag was transfected simultaneously. After 24 h, the cells were infected with IME-BJ01 strains. The mRNA levels of M and E genes of IME-BJ01 strain in the cells (A, D) and E gene in supernatant (B, E) were detected by RT-qPCR, as well as the detection of viral titers in supernatant (C, F). The N protein of the virus and the overexpression efficiency was detected by IB (G, H). Data are representative of three independent experiments and shown as average ±SD (n = 3). Significance was determined by one-way ANOVA, followed by a Tukey multiple comparisons posttest. P > 0.05; **P < 0.01; ***P < 0.001. Figure 6—figure supplement 1 shows data related to infection with Omicron BA.1. Figure 6-source data 1. PDF file containing original western blots for Figure 6G and H, indicating the relevant bands and treatments. Figure 6-source data 2. Original files for western blot analysis displayed in Figure 6G and H. Figure 6-source data 3. Numerical data obtained during experiments represented in Figure 6.

### UBR5 and MARCHF7 mediate nsp16 variant degradation

With the continuous emergence of SARS-CoV-2 variants, broad-spectrum antiviral strategies have become essential. To assess whether UBR5 and MARCHF7 mediate various nsp16 variants, we analysed nsp16 amino acid sequences from the National Center for Biotechnology Information (NCBI) (Figure 6—figure supplement 3A). Most variants had one or two mutations, except for XBB.1.9.1, XBB.1.5, and XBB.1.16, which contained more mutations, suggesting a conserved sequence. We used mutagenesis to synthesise nsp16 sequences for these three variants, along with several single-site mutants from other variants. Treatment with MG132 restored nsp16 protein levels for all variants (Figure 6—figure supplement 3B), confirming UPS-mediated degradation. Furthermore, *UBR5* or *MARCHF7* knockdown stabilised nsp16 proteins from these variants, though to varying degrees (Figure 6—figure supplement 3C). These results suggest that UBR5 and MARCHF7 contribute to broad-spectrum antiviral activity by targeting nsp16 from diverse SARS-CoV-2 variants for degradation.

### SARS-CoV-2 infection reduces UBR5 and MARCHF7 expression

To investigate the impact of SARS-CoV-2 infection on UBR5 and MARCHF7 expression, we analysed their mRNA and protein levels following infection with the IME-BJ01 and Omicron strains at varying MOIs using RT-qPCR and immunoblotting. MARCHF7 expression decreased consistently with increasing viral titres, whereas UBR5 levels initially rose at low titres before declining as infection progressed (Figure 6—figure supplement 4A-4C). This transient UBR5 upregulation may be linked to its role in interferon-γ-mediated pathways, as previously reported (Wu et al., 2022). To confirm these findings in vivo, we examined UBR5 and MARCHF7 mRNA levels in peripheral blood mononuclear cells from SARS-CoV-2-infected patients with varying disease severity. UBR5 expression negatively correlated with disease progression, whereas MARCHF7 levels showed no significant correlation (Figure 6—figure supplement 4D). These results suggest that SARS-CoV-2 may actively suppress host antiviral defences by downregulating UBR5 and MARCHF7 expression.

### UBR5 and MARCHF7 protect mice from SARS-CoV-2 challenge

Since no specific activators or inhibitors for E3 ligases are available, we used a transient overexpression method using high-pressure tail vein injection of plasmids in mice (Bonamassa et al., 2011). To evaluate their antiviral effects in vivo, mice were injected with plasmids encoding UBR5 or MARCHF7, followed by SARS-CoV-2 challenge (Fig. 7A). E3 ligase overexpression significantly reduced viral E gene copy numbers in the lungs and markedly decreased viral titres (Fig. 7C-D). Additionally, treated mice experienced less weight loss compared to controls (Fig. 7E). Histopathological analysis at 5 days post-infection showed that lungs from UBR5- or MARCHF7-treated mice exhibited reduced alveolar contraction and pulmonary oedema compared to the severe lesions observed in control mice (Fig. 7F). Immunoblotting further confirmed lower N protein abundance in treated mice (Fig. 7G). These results suggest that UBR5 and MARCHF7 overexpression suppresses SARS-CoV-2 virulence in vivo by promoting nsp16 degradation.

**Figure 7.**
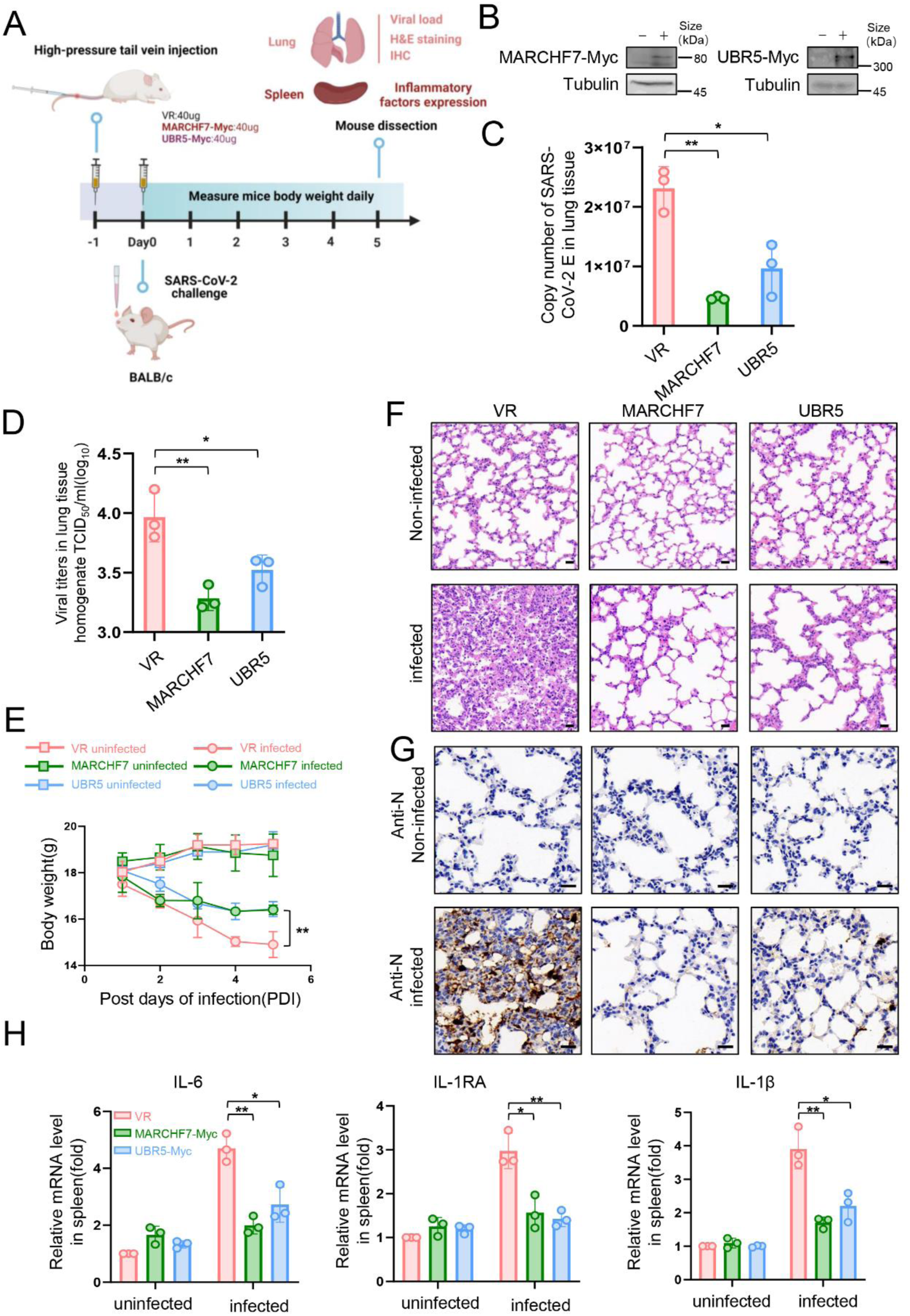
In a mouse infection model, overexpression of MARCHF7 or UBR5 exerted inhibitory effects on virus. (A-G) BLAB/C mice were injected with the corresponding plasmids at 40ug/500ul via the high-pressure tail vein, followed by nasal inoculation with 50µl SARS-CoV-2 virus at a dosage of 10^5.5^ TCID50/mL (created with BioRender.com and the agreement no. is FO27O5X39A). IB was used to detect the expression of MARCHF7 or UBR5 in the lung tissues (B). Viral RNA loads in mouse lung tissues were detected by measuring the mRNA levels of the E genes by RT-qPCR (C). Lung tissue was collected, homogenized, and the residue was removed by centrifugation to collect the supernatant. The viral titer was then measured using the TCID50 method (D). Mouse body weight was monitored during the experimental period (E). Representative images of H&E staining of lungs of mice with different treatments. Magnification, ×40. Bars, 20 µm (F). The staining of viral N proteins. Magnification, ×63. Bars, 20 µm. n =3 in each group (G). RT-qPCR was used to measure the expression of cytokines and chemokines in the spleens of mice in each group (H). Statistical significance was analyzed using a one-way analysis of variance with Tukey’s multiple comparisons test. (NS, no significance, *p < 0.05, **p < 0.01, ***p < 0.001). Figure 7-source data 1. PDF file containing original western blots for Figure 7B, indicating the relevant bands and treatments. Figure 7-source data 2. Original files for western blot analysis displayed in Figure 7B. Figure 7-source data 3. Numerical data obtained during experiments represented in Figure 7.

**Figure 8.**
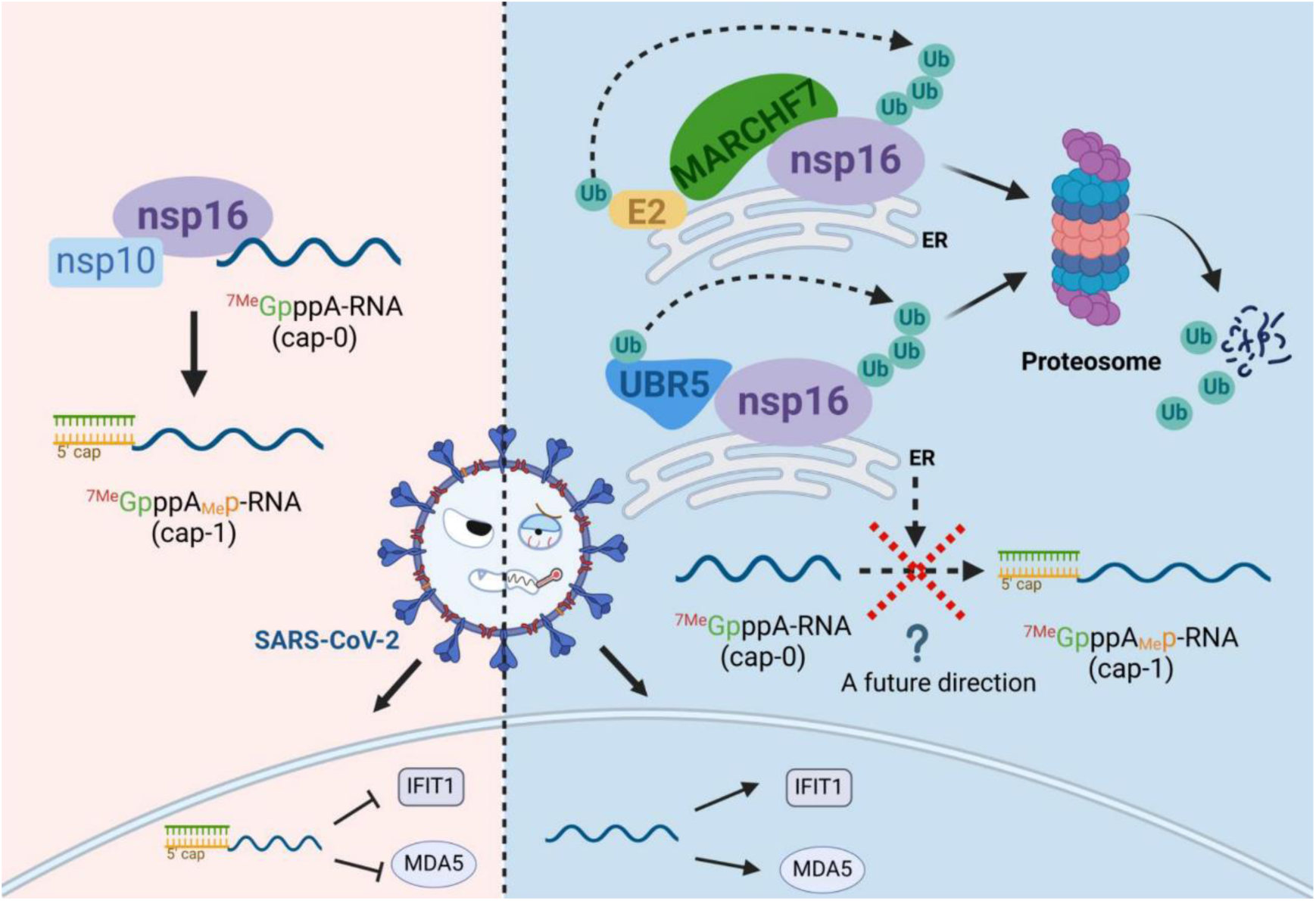
Schematic diagram of MARCHF7 and UBR5 ubiquitinate the SARS-CoV-2 non-structural protein nsp16, leading to its degradation via the proteasomal pathway, thereby affecting viral replication (created with BioRender.com and the agreement no. is FM27O5ZS3U).

SARS-CoV-2 infection can induce a severe cytokine storm, contributing to high mortality rates (Song et al., 2020) (Hojyo et al., 2020). To assess the impact of UBR5 and MARCHF7 on inflammatory responses, we measured key cytokine levels—including interleukin-6 (IL-6), interleukin-1 receptor antagonist (IL-1RA), and interleukin-1β (IL-1β)—in the spleens of infected mice (Makaremi et al., 2022) (Hu et al., 2021). Consistent with their antiviral activity, UBR5 and MARCHF7 treatment significantly reduced the production of these inflammatory cytokines (Fig. 7H).

## Discussion

SARS-CoV-2 nsp16 functions as a 2′-O-methyltransferase (2′-O-MTase), catalysing penultimate nucleotide methylation of the viral RNA cap. This modification allows SARS-CoV-2 to mimic host mRNA, evading immune detection and response (Lin et al., 2020) (Balieiro et al., 2022; Russ et al., 2022). High-resolution structures of nsp16 and nsp16-nsp10 heterodimers have been characterised (Rosas-Lemus et al., 2020) (Klima et al., 2022; Lugari et al., 2010), providing a foundation for antiviral drug development targeting these complexes (Balieiro et al., 2022; Melo-Filho et al., 2022). However, host factors that interact with and regulate nsp16 remain largely unknown.

Previous studies, including work from our group, have highlighted the role of the UPS in modulating SARS-CoV-2 infection (Xu et al., 2022). Several host E3 ligases—such as RNF5, Cullin4-DDB1-PRPF19, ZNF598, and TRIM7—ubiquitinate and degrade key SARS-CoV-2 proteins, thereby suppressing viral replication (Li et al., 2023; Liang et al., 2022; Maimaitiyiming et al., 2022; Zhang et al., 2024). In contrast, SARS-CoV-2 uses deubiquitinases (DUBs) to counteract these defences and enhance viral replication (Chen et al., 2024; Gao et al., 2024; Gao et al., 2022; Guo et al., 2021). In this study, we identify nsp16 as a short-lived protein with a half-life of only 2 – 3 h, which is targeted for ubiquitination and degradation via the UPS. Using MS and bioinformatics screening, we identified two E3 ligases, UBR5 and MARCHF7, that directly interact with nsp16 and mediate nsp16 ubiquitination and degradation.

UBR5, a member of the UBR box protein family, contains a HECT domain essential for its E3 ligase activity (Kim et al., 2021). UBR5 regulates tumour growth, metastasis (Xiang et al., 2022) (Qiao et al., 2020), and viral infections. For instance, UBR5 targets the ORF4b protein of Middle East respiratory syndrome coronavirus (MERS-CoV) for degradation, reducing its ability to counteract the host immune response (Zhou et al., 2022). Conversely, UBR5 supports Zika virus replication by assisting TER94/VCP in degrading the capsid protein, allowing viral genome release into the cytoplasm (Gestuveo et al.). MARCHF7, a RING E3 ligase, plays a role in T cell proliferation, neuronal development, inflammasome regulation, and tumour progression (Zheng, 2021) (Zhang et al., 2016) (Cai et al., 2022) (Zhao et al., 2018). However, its involvement in viral infections remains largely unexplored, with one study identifying it as a host protein interacting with the 2C protein of foot-and-mouth disease virus (Mahajan et al., 2021).

Our findings demonstrate that UBR5 and MARCHF7 independently interact with nsp16 and facilitate its degradation, primarily on the cytoplasmic ER. Our findings highlight the ongoing evolutionary battle between viruses, which evolve strategies to evade host defences, and hosts, which deploy multiple mechanisms to counteract viral infections. One such strategy is the use of multiple E3 ligases targeting the same viral protein for degradation. Functional analyses revealed that knocking down *UBR5* or *MARCHF7* enhanced SARS-CoV-2 replication, whereas their overexpression impaired it. Moreover, when their enzymatic activity was inactivated, the antiviral effect was abolished, confirming that their E3 ligase activity is essential for viral suppression. Notably, nsp16 overexpression alone did not increase viral levels in the absence of functional UBR5 or MARCHF7. This underscores the critical antiviral role of UBR5 and MARCHF7 in restricting SARS-CoV-2 by promoting nsp16 degradation.

Due to the presence of 18 lysine residues in nsp16, identifying specific ubiquitination sites was challenging. Using structural data (Decroly et al., 2011), we constructed truncated nsp16 mutants to assess their degradation profiles. Our results showed that nsp16 mutants degradation occurred to varying degrees (Figure 8—figure supplement 1A-B), suggesting that multiple lysine residues serve as recognition sites for E3 ligases. Furthermore, MG132 recovery experiments provided valuable insights. The Δ2-17 mutant, which lacks this segment, exhibited more than a two-fold increase in nsp16 expression, whereas the Δ204-212 mutant showed significantly reduced recovery after MG132 treatment compared to the wild-type. These observations offer important clues for further investigation into the interaction regions between nsp16 and MARCHF7 or UBR5, as well as potential ubiquitination sites. MS analysis identified K76 as a ubiquitination site (Figure 8—figure supplement 1C). However, the nsp16 K76R mutant still underwent degradation, though its ubiquitination level was reduced compared to that of the wild-type, indicating that K76 is one of several ubiquitination sites (Figure 8—figure supplement 1D-E). This suggests that nsp16 may undergo ubiquitination at multiple sites or through non-canonical pathways at non-lysine residues (McClellan et al., 2019). Beyond UBR5 and MARCHF7, other E3 ligases may also contribute to nsp16 regulation, warranting further investigation into additional host factors involved in SARS-CoV-2 restriction.

Currently, no specific activators or inhibitors exist for UBR5 or MARCHF7. To assess their antiviral effects in vivo, we used high-pressure tail vein injection, a gene delivery method primarily targeting the liver (Kamimura et al., 2014) but also effective for lung tissues (Bonamassa et al., 2011). Gene expression was successfully detected in the lungs, and while this method induced mild immune activation, its effect was minimal compared to other techniques (Suda et al., 2023). Notably, mice and non-human primates exhibit greater resistance to innate immune activation than humans (Raper et al., 2003), suggesting that immune activation had little influence on our results. However, translating these findings into therapeutic applications remains challenging. For instance, human trials using naked DNA injections (e.g., OTC cDNA) have shown variable immune responses, with one case of fatal immune activation reported (Raper et al., 2003). This underscores the variability of human immune responses and the challenges in translating findings from animal models to clinical applications. Such variability may also explain the inconsistent trends in MARCHF7 mRNA levels observed in peripheral blood samples from patients with different disease severities. Additionally, the small sample size in our study may have contributed to this observation.

In conclusion, in this study, we identified the host E3 ligases UBR5 and MARCHF7 as key regulators of SARS-CoV-2 nsp16, facilitating its ubiquitination and degradation. We observed a negative correlation between their expression levels and infection severity, highlighting their antiviral function. Moreover, both ligases effectively suppressed SARS-CoV-2 replication. These findings provide a foundation for developing UPS-targeted therapeutic strategies for COVID-19 treatment.

## Materials and Methods

A list of materials is partially provided in Supplementary file 1.

### Reagents and antibodies

The drugs used in this study were as follows: MG132 (catalog no. S2619), Bafilomycin A1 (catalog no. S1413), Bortezomib (catalog no. S1013), Carfilzomib (catalog no. S2853) and Vinblastine (catalog no. S4504) were purchased from Selleck (Houston, TX, USA). Cycloheximide (catalog no. 66-81-9) was purchased from Sigma (Saint Louis, MO, USA). The antibodies used for immunoblotting analysis (IB) were as follows: anti-UBR5 mAb (Proteintech, Rosemont, IL, USA, catalog no. 66937-1-Ig, RRID: AB_2881136), anti-MARCHF7 mAb (Santa Cruz Biotechnology, Dallas, TX, USA, catalog no. sc-166945, RRID: AB_2157974), SARS-CoV-2 2’-O-ribose Methytransferase Antibody (nsp16) (Cell signaling, Danvers, Massachusetts, USA, catalog no. #70811, RRID: AB_2799787), anti-Flag mAb (Sigma, catalog no. F1804, RRID: AB_262044), anti-tubulin mAb (Abcam, Cambridge, Cambridgeshire, UK, catalog no. ab11323, RRID: AB_297919), anti-hemagglutinin (anti-HA) pAb (Invitrogen, catalog no. 71-5500, RRID: AB_2533987), anti-Myc pAb (Proteintech, Rosemont, IL, USA, catalog no. 16286-1-AP, RRID: AB_2877901), and anti-GFP mAb (Abcam, catalog no. ab1218, RRID: AB_298079). Primary antibody: anti-COX5A pAb (Sangon Biotec, Shanghai, CHN, catalog no. D261450), anti-PDI-mAb (Proteintech, catalog no. 2E6A11, RRID: AB_2881137) and Human GM130/GOLGA2 Antibody (R&D Systems, Minneapolis, MN, USA, catalog no. AF8199, RRID: AB_2136393) were used for immunofluorescence, and SARS-CoV-2 nucleocapsid monoclonal antibody (mAb) (GeneTex, Irvine, CA, USA, catalog no. GTX635679, RRID: AB_2909916) were used for Immunohistochemistry. Fluorescent secondary antibodies: goat anti-Rabbit IgG (H+L) Highly Cross Adsorbed Secondary Antibody, Alexa Fluor Plus 488 (Invitrogen, catalog no. A11001, RRID: AB_2762824), and goat anti-Rabbit IgG (H+L) Highly Cross Adsorbed Secondary Antibody, Alexa Fluor Plus 568 (catalog no. A-11011, RRID: AB_143157) and immunohistochemical secondary antibodies: Streptavidin-Peroxidase Anti-Rabbit IgG kit (catalog no. KIT-9706) were purchased from Invitrogen and maixin (Fuzhou, CHN) respectively.

### Cell lines and viruses

The cell lines used in this study include HEK293T cells (catalog no. CRL-11268), Hela cells (catalog no.CRM-CCL-2), Caco2 cells (catalog no. HTB-37). All cell lines were purchased from American Type Culture Collection (ATCC; Manassas, VA, USA) and tested for mycoplasma. In order to conduct virus related experiments, stable cell lines including Caco2-N^int^ (stably expressing SARS-CoV-2 N gene) and 293T-ACE2 (stably expressing ACE2) were generated in our laboratory. The cells were cultured in a cell incubator containing 5% CO2 at 37°C using Dulbecco’s modified Eagle’s medium (DMEM; HyClone, Logan, UT, USA) containing 10% heat-inactivated fetal calf serum (FCS, GIBCO BRL, Grand Island, NY, USA), 100 mg/ml penicillin, and 100 ug/ml streptomycin to provide nutrition. We expanded SARS-CoV-2 virus-like-particles (GenBank access no. MN908947, Clade:19A) with high replication capacity in Caco2-N^int^ using a biosafety level-2 (BSL-2) cell culture system. The virus infecting the cells include Omicron BA. 1 strain (human/CHN_CVRI-01/2022) and SARS-CoV-2 IME-BJ01 strain (BetaCoV/Beijing/IME-BJ05-2020, GenBank access no. MT291831.1), and mouse adapted SARS-CoV-2/C57MA14 variant (GenBank: OL913104.1) used for experiments in mice were provided by Key Laboratory of Jilin Province for Zoonosis Prevention and Control, and all experiments for virus infection were performed in Biosafety level 3 (BSL-3) cell culture system.

### Plasmids

Eukaryotic expression plasmids encoding SARS-CoV-2 nsp protein were provided by Professor Wang Peihui (Shandong University). The nsp16-HA expression vector was constructed by adding HA tag at the C-terminus. UBR5 (Gnen ID: 51366) and its mutants (UBR5-ΔHECT, UBR5-ΔPABC, UBR5-ΔUBR) expression plasmids were constructed using purchased plasmids from Addgene (Watertown, MA, USA) as templates with no tag or a MYC tag at the N-terminus. The cDNA of 293T cells was used as a template to construct MARCHF7(Gene ID: 64844) and its truncated mutant with a Myc tag at the C-terminus. All the above expression plasmids were inserted into the VR1012 vector. For immunofluorescence (IF) and Fluorescence Resonance Energy Transfer (FRET) experiments, pCDNA3.1-YFP vector (catalog no. 13033) and pECFP-C1 vector (catalog no. 6076-1) were purchased from Addgene and BD (Biosciences Clontech), and MARCHF7 or UBR5 was constructed on pECFP-C1, and nsp16 was constructed on pCDNA3.1-YFP. Single point mutants of nsp16 of different virus subtypes were obtained by point mutagenesis. Multi-site combined mutants were synthesized by Sangon Biotec Company. Human ubiquitin protein and its mutants have been previously described. Primers required for PCR were listed in Supplementary file 1.

### RNA extraction and real-time quantitative PCR

We used the way of Trizol (Invitrogen, catalog no. 15596018CN) to extract RNA. The RNA was subsequently reverse transcribed using a High-Capacity cDNA Reverse Transcription kit (Applied Biosystems, Carlsbad, CA, USA, catalog no. 4368814). Finally, PrimeScript RT Master Mix (Takara, Shiga, JPN, catalog no. RR036A) and relative real-time primer were used for qPCR on an Mx3005P instrument (Agilent Technologies, Stratagene, La Jolla, CA, USA). Amplification procedure of the target fragment is as follows: predenaturation at 95°C for 2min, denaturation at 95 °C for 30s, annealing at 55 °C for 30s, and extension at 72 °C for 30s, total of 40 cycles. Real-time primer sequences used in this study were shown in Supplementary file 1.

### Immunoblotting analysis (IB)

Cells or supernatant cultured for a certain time were collected, cell precipitates were resuspended with lysis buffer (50 mM Tris–HCl [pH 7.8], 150 mM NaCl, 1.0% NP-40, 5% glycerol and 4 mM EDTA), 4xloading buffer was added (0.08M Tris [pH 6.8], 2.0%SDS, 10% glycerol, 0.1 M dithiothreitol, and 0.2% bromophenol blue), and samples were lysed by heating at 100℃ for 30min. After removal of cell debris by centrifugation, the supernatant was taken to separate proteins of different sizes by SDS-PAGE. The proteins were transferred onto polyvinylidene fluoride (PVDF) membranes, incubated overnight with the indicated primary antibody, and after 1 hour at room temperature with HRP-conjugated secondary antibodies (Jackson Immunoresearch, West Grove, NJ, USA, catalog no. 115-035-062 for anti-mouse and 111-035-045 for anti-rabbit), the proteins were visualized through Ultrasensitive ECL Chemiluminescence Detection Kit (Proteintech, catalog no. B500024).

### Stable cell line generation

We purchased the lentiviral vector pLKO.1-puro (catalog no. 8453) from Adggen. ShRNA targeting target genes were designed and synthesized by Sangon Biotech. After annealing, it was inserted into the vector. Lentivirus was generated by transfection with shRNA (cloned in pLKO.1) or control vector, RRE, VSVG and REV, and the cells were screened with 1/1000 puromycin (3 μg/ml, Sigma, catalog no. P8833) after infection 48 hours. The knockdown efficiency was detected by RT-qPCR or IB. The shRNA target sequences used in this study are shown in Supplementary file 1.

### RNAi

Short interfering RNA (siRNA) used in this study were purchased from RiboBio Co. Ltd. (Guangzhou, CHN). The siRNA sequences for knockdown of *MARCHF7* or *UBR5* are provided in Supplementary file 1. The siRNA was transfected into the cells by transfection reagent Lipofectamine 2000 (Invitrogen, catalog no. 11668-019), and the corresponding plasmids were transfected 24 hours later using Lipofectamine 3000 Reagent (Invitrogen, Carlsbad, CA, USA, catalog no. L3000-008). The expression of target genes was detected by RT-qPCR or immunoblotting (IB) analysis 48 hours later.

### Immunoprecipitation

After 10 hours of MG132 (10 µM) treatment, the cells were harvested, resuspended in 1ml lysis buffer (50 mM Tris-HCl [pH 7.5], 150 mM NaCl, 0.5% NP-40) containing protease inhibitor (Roche, catalog no. 11697498001), placed on the shaker oscillator for 4 hours to lyse, then centrifuged to remove cell debris, and the supernatant was incubated with the antibody and protein G agarose beads (Roche, Basel, Basel-City, CH, catalog no. 11243233001) at 4°C overnight. Protein G agarose beads was washed 6-8 times with wash buffer (20 mM Tris-HCL [pH 7.5], 100 mM NaCl, 0.1 mM EDTA, and 0.05% Tween 20) at 4°C, centrifuged at 800 g for 1 min. The proteins were eluted by adding loading buffer (0.08 M Tris [pH 6.8], 2.0% SDS, 10% glycerol, 0.1 M dithiothreitol, and 0.2% bromophenol blue), the samples were boiled at 100°C for 10min to elate the proteins. The lysates and immunoprecipitations were detected by IB.

### Mass spectrometry (MS)

HEK293T were transfected with SARS-CoV-2-nsp16-Flag for 36 hours, and then harvested after 10 hours of treatment with MG132 or DMSO. Proteins were enriched by co-immunoprecipitation assay. And the elutions were analyzed by MS. MS analysis was performed by the national center for protein science (Beijing, CHN).

### Immunofluorescence

At 48 hours after transfection, the solution was discarded and cells were washed twice with preheated PBS for 5min each, followed by fixation with 4% paraformaldehyde at 37°C for 10min. After three washes with PBS, permeabilized with 0.2% Triton X-100 for 5 min at 37°C, and then immediately followed by blocking with 10% fetal bovine serum at room temperature for 1 hour. The blocked cells were incubated overnight with the indicated primary antibodies. The next day, after three washes with PBS, the cells were incubated with corresponding secondary antibodies for 1 hour at room temperature in the dark. The nuclei were stained with DAPI (49,6-diamidino-2-phenylindole, Sigma, catalog no. 9542) and then stored in 90% glycerol. The fluorescence was detected by a laser scanning confocal microscope (FV 3000, Olympus, Tokyo, Japan).

### Viral infectivity assay

The Caco2 cells were transfected with siRNA by Lipofectamine RNAiMAX Reagent (Invitrogen, catalog no. 13778150) to knockdown *UBR5* or *MARCHF7* and infected with IME-BJ01 strain (MOI=0.01) or Omicron-BA.1 strain (MOI=0.001) at 24 hours after transfection. After two hours of infection, the Caco2 cells were cultured in fresh medium containing 2% fetal bovine serum for 48 hours. The cells and supernatants were collected. Viral levels were characterized by RT-qPCR and IB analysis of viral structural proteins. The overexpression plasmid UBR5-Myc, MARCHF7-Myc and nsp16-Flag was transfected in 293T-ACE2 by Lipofectamine 3000 Reagent and infected with virus 24 hours later.

### Mouse lines and infections

BALB/C mice, 6 weeks, were purchased from Charles River Laboratories, Beijing. Mice were cultured in groups according to experimental groups. Our study examined male and female animals, and similar findings are reported for both sexes. To reduce animal suffering, all welfare and experimental procedures were carried out in strict accordance with the Guide for the Care and Use of Laboratory Animals and relevant ethical regulations during the experiment. The mice were randomly divided into 6 groups and injected with 40 µg/500 µl MARCHF7 or UBR5 plasmids via the tail vein at high pressure. Three groups of mice were infected 50ul with SARS-CoV-2 isolates by nasal challenge at a dose of 10^5.5^ TCID_50_/mL, three groups were not infected with virus as control, and the empty vector injection group was also used as control.

### Immunohistochemical analysis

Three mice in each group, a total of 18 mice, were sacrificed after anesthesia. The lungs of mice were collected and fixed in 4% paraformaldehyde fixative. After 2 days, the lungs were progressively dehydrated through different concentrations of ethanol solution, transparently treated, soaked in xylene, and finally embedded in paraffin. Some thin sections obtained through a microtome for hematoxylin and eosin (H&E) staining. The other fraction was incubated with 3% hydrogen peroxide for 5-10 min at room temperature to eliminate endogenous peroxidase activity. SARS-CoV-2 nucleocapsid monoclonal antibody (mAb) (GeneTex, catalog no. GTX635679) and a Streptavidin-Peroxidase Anti-Rabbit IgG kit (Maixin, Fuzhou, CHN, catalog no. KIT-9706) was used to quantify the viral level in lung tissue.

### Statistical analysis

The statistical analyses used in the figures have been described in detail in the figure legends. All data were expressed as mean ± standard deviations (SDs). Statistical comparisons were performed using student’s t-test, one-way ANOVA, or repeated measurement ANOVA. The differences were statistically significant as follows: *P < 0.05, **P < 0.01, ***P < 0.001; ns is for no meaning

### Ethics statement

Collection of inpatient blood was approved by the Ethics Committee of the First Hospital of Jilin University (21K105-001) in accordance with the guidelines and principles of The World Medical Association (WMA) Declaration of Helsinki and the Department of Health and Human Services Belmont Report. All study participants signed an informed consent form. PBMCS were obtained from 9 COVID-19 patients with mild disease, 6 patients with severe disease, and 5 patients with critical disease (Supplementary file 2). All animal experiments were approved by the ethics committee of Research Unit of Key Technologies for Prevention and Control of Virus Zoonoses, Chinese Academy of Medical Sciences, Changchun Veterinary Research Institute, Chineses Academy of Agricultural Sciences (IACUC of AMMS-11-2020-006).

## Acknowledgments

We thank C.Y. Dai for providing critical reagents.

## Funding information

This work was supported by funding from the National Natural Science Foundation of China (82341072 and 82272316 to ZWY, 82341062 to LZL), the National Key R&D Program of China (2021YFC2301900 and 2301904, 2023YFC2306603), the Science and Technology Department of Jilin Province (YDZJ202301ZYTS521 and YDZJ202201ZYTS587), the Key Laboratory of Molecular Virology, Jilin Province (20102209), and Bethune Project, Jilin University (2023B03). The funding sources were involved in study design, data collection and interpretation, and the decision to submit the work for publication.

## Data availability

The authors declare that all data supporting the findings of this study are available within the paper and its supplementary information files. The mass spectrometry proteomics data have been deposited to the ProteomeXchange Consortium (http://proteomecentral.proteomexchange.org) via the iProX partner repository with the dataset identified PXD053961.

## Figure legends

**Figure 1—figure supplement 1.**
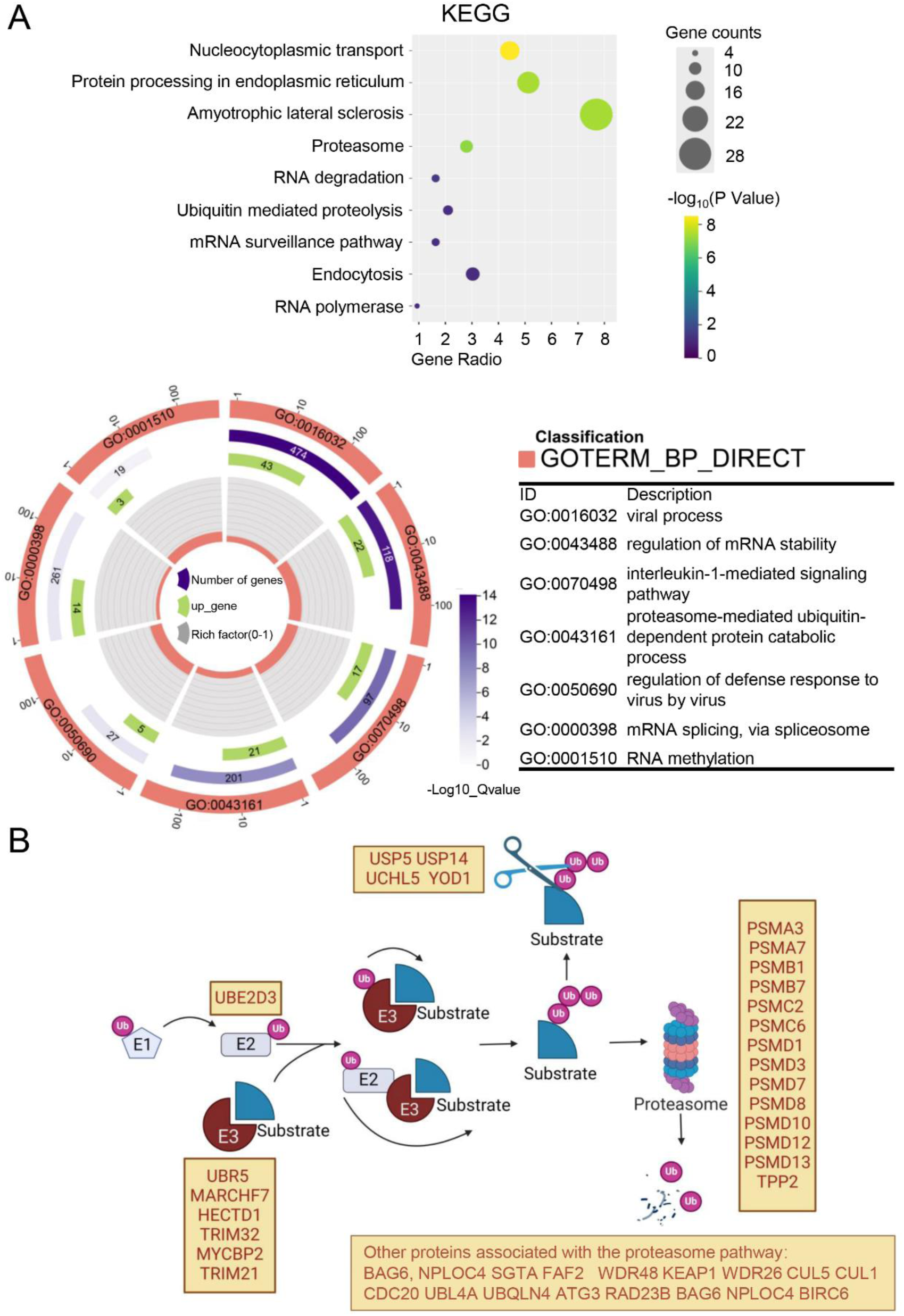
Analysis of mass spectrometry results. A. The DAVID bioinformatics website (https://david.ncifcrf.gov/tools.jsp) was used for KEGG pathway analysis and GO analysis of the differential proteins bound by nsp16 protein after treating with or without MG132. The bubble plots or circle diagram depict the results of the analysis separately. In bubble plots, a change from yellow to purple indicates a decreased p-value, while the size of the circles indicates the number of enriched genes. In the circle plot, depicting the BP (Biological process) analysis in the GO term, the length of the purple rectangle indicates the number of genes included in the term. The length of the green rectangle indicates the number of overlapping genes between the genes included in the term and the genes entered in the gene enrichment analysis, a change from deep to shallow of purple indicates a decreased p-value (created with chiplot.com). B. Schematic representation of proteins degraded by the proteasome pathway, as well as proteins associated with the proteasome in the MS enrichment analysis are shown (created with BioRender.com and the agreement no. is VO27O5XHHC).

**Figure 2—figure supplement 1.**
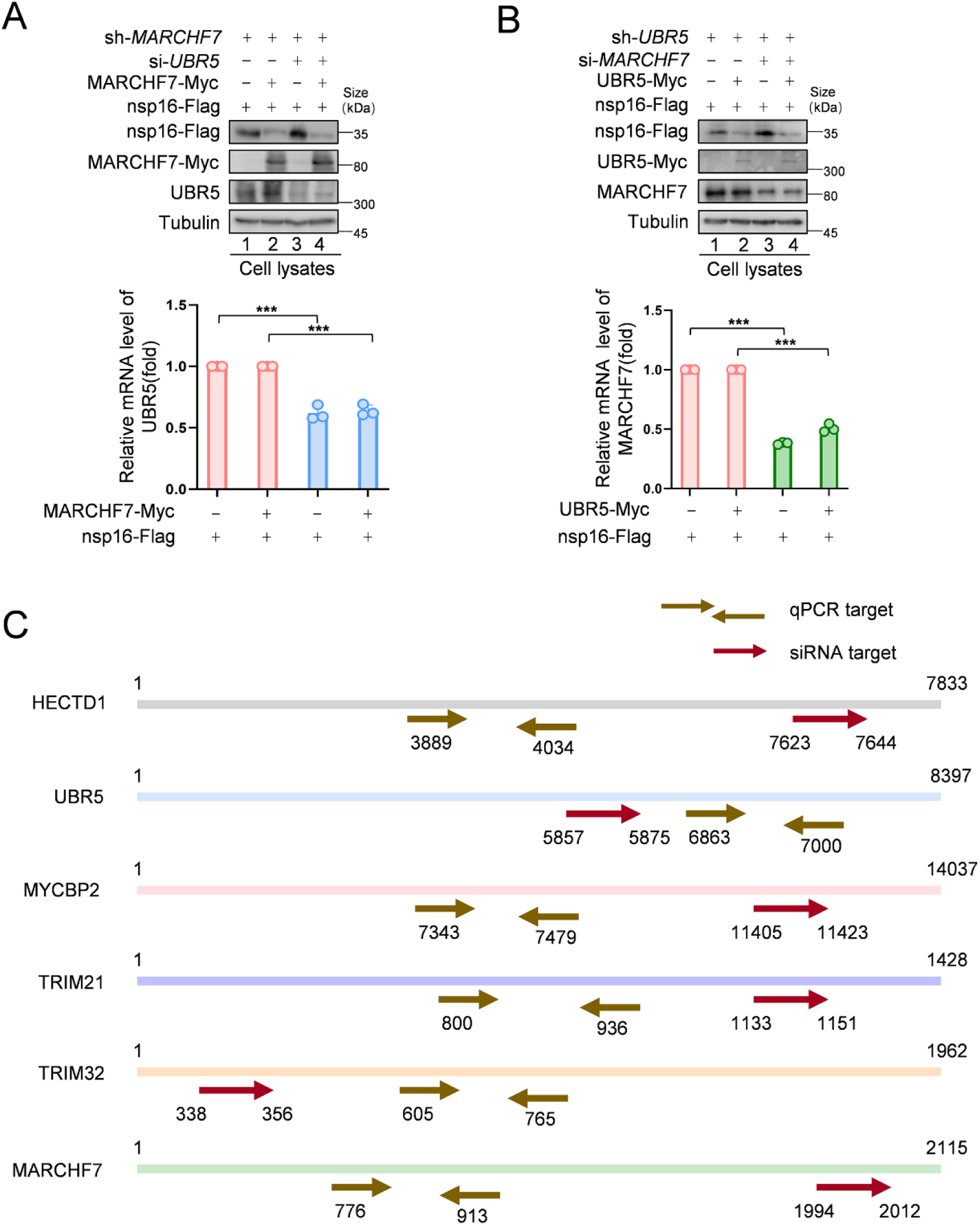
MARCHF7 and UBR5 degrade nsp16 independently. (A-B). *UBR5* siRNA was transfected into shMARCHF7 cells to knock down *UBR5*. After 24 hours, MARCHF7 and nsp16 expression vectors were co-transfected, and the cells were harvested 72 hours later. The levels of nsp16 were characterized by IB with anti-Flag antibody. Whether MARCHF7 was dependent on UBR5 to degrade of nsp16 was determined by further transfection of *MARCHF7* siRNA into shUBR5 cells, followed by co-transfection of UBR5 and nsp16 expression vectors 24 h later. The other operations are the same as above. Data are representative of three independent experiments and shown as average ±SD (n = 3). Significance was determined by a two-tailed t-test. *P > 0.05; **P < 0.01; ***P < 0.001. A. The siRNA targeting regions and RT-qPCR targeting regions for the E3 ubiquitin ligases: HECTD1, UBR5, MYCBP2, TRIM21, TRIM32, and MARCHF7 are shown. Figure 2—figure supplement 1-source data 1. PDF file containing original western blots for Figure 2—figure supplement 1A and B, indicating the relevant bands and treatments. Figure 2 — figure supplement 1-source data 2. Original files for western blot analysis displayed in Figure 2—figure supplement 1A and B. Figure 2—figure supplement 1-source data 3. Numerical data obtained during experiments represented in Figure 2—figure supplement 1.

**Figure 4—figure supplement 1.**
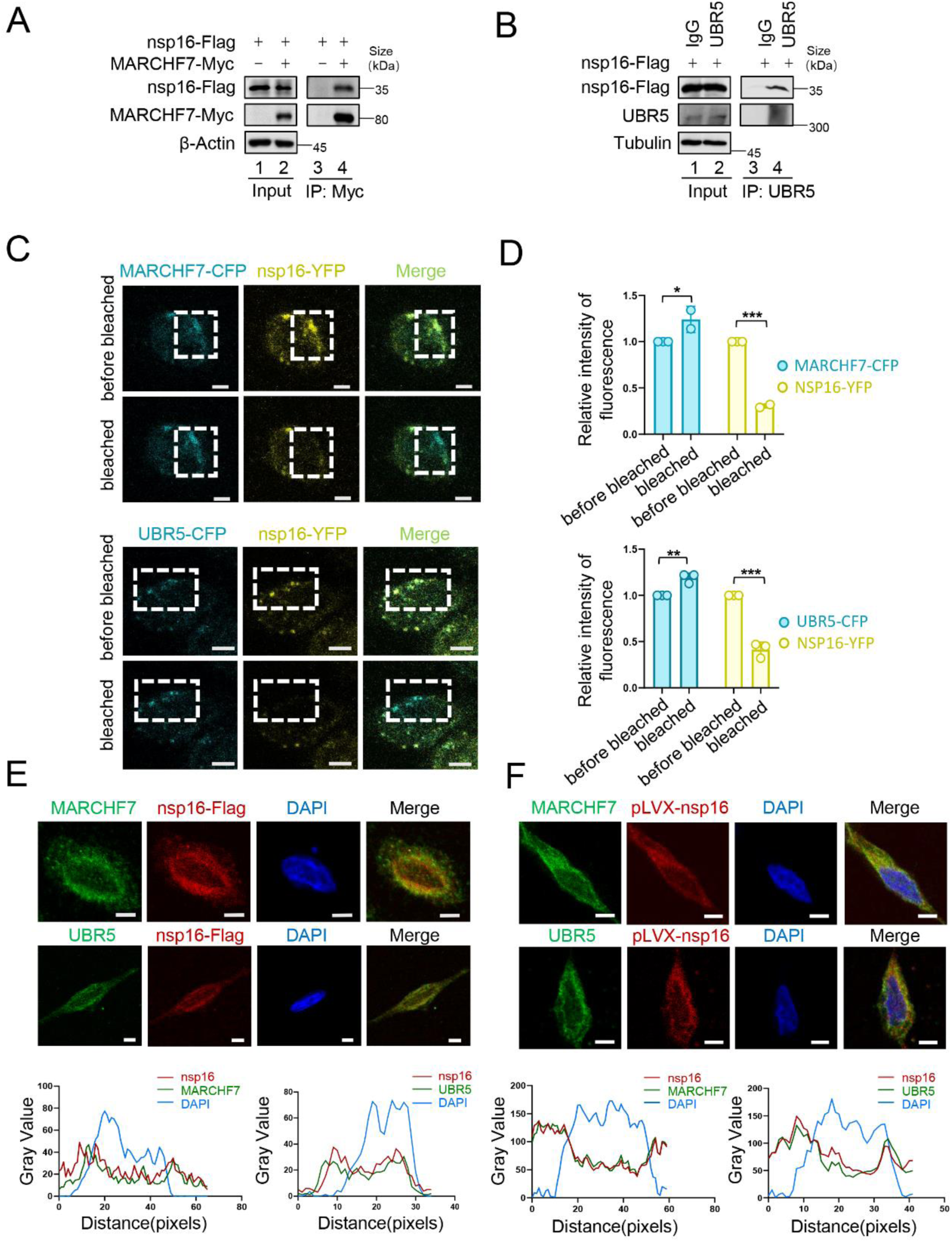
Interaction of MARCHF7 or UBR5 with nsp16. A. HEK293T cells were transfected with either nsp16-Flag alone or together with MARCHF7-Myc. Thirty-six hours after transfection, the cells were treated with MG132 (10 µM) for 12 h. Cell lysates were subjected to immunoprecipitation with anti-Flag antibody. Using IB to analyze the precipitates and input. B. HEK293T cells were transfected with nsp16-Flag. Cell lysates were subjected to immunoprecipitation with anti-UBR5 or IgG antibody. (C-D). Hela cells were co-transfected with YFP-nsp16 and CFP-UBR5 or CFP-MARCHF7. A representative image of YFP-nsp16 (yellow) and ECFP-MARCHF7 (cyan) or ECFP-UBR5 (cyan) expressing cells before and after photobleaching the acceptor fluorophore, YFP. The region chosen for photobleaching is marked (white open box), Bars, 10um. The quantization of fluorescence brightness was analyzed by Image J. Data are representative of three independent experiments and shown as average ±SD (n = 3). Significance was determined by a two-tailed t-test. *P > 0.05; **P < 0.01; ***P < 0.001. E. HeLa cells transfected with nsp16-Flag were analyzed by confocal microscopy. The Flag-tagged nsp16 labeled with anti-Flag antibody (red). MARCHF7 or UBR5 were labeled with endogenous antibodies (green). Cell nuclei were stained using DAPI (blue). Representative images were shown. Scale bars, 20 um. The ratio of colocalization was quantified by measuring the fluorescence intensities using Image J. F. nsp16 was stably transfected into HEK293T cells. The cells were analyzed by confocal microscopy. The other operations are the same as above. Figure 4—figure supplement 1-source data 1. PDF file containing original western blots for Figure 4—figure supplement 1A and B, indicating the relevant bands and treatments. Figure 4 — figure supplement 1-source data 2. Original files for western blot analysis displayed in Figure 4—figure supplement 1A and B. Figure 4—figure supplement 1-source data 3. Numerical data obtained during experiments represented in Figure 4—figure supplement 1.

**Figure 4—figure supplement 2.**
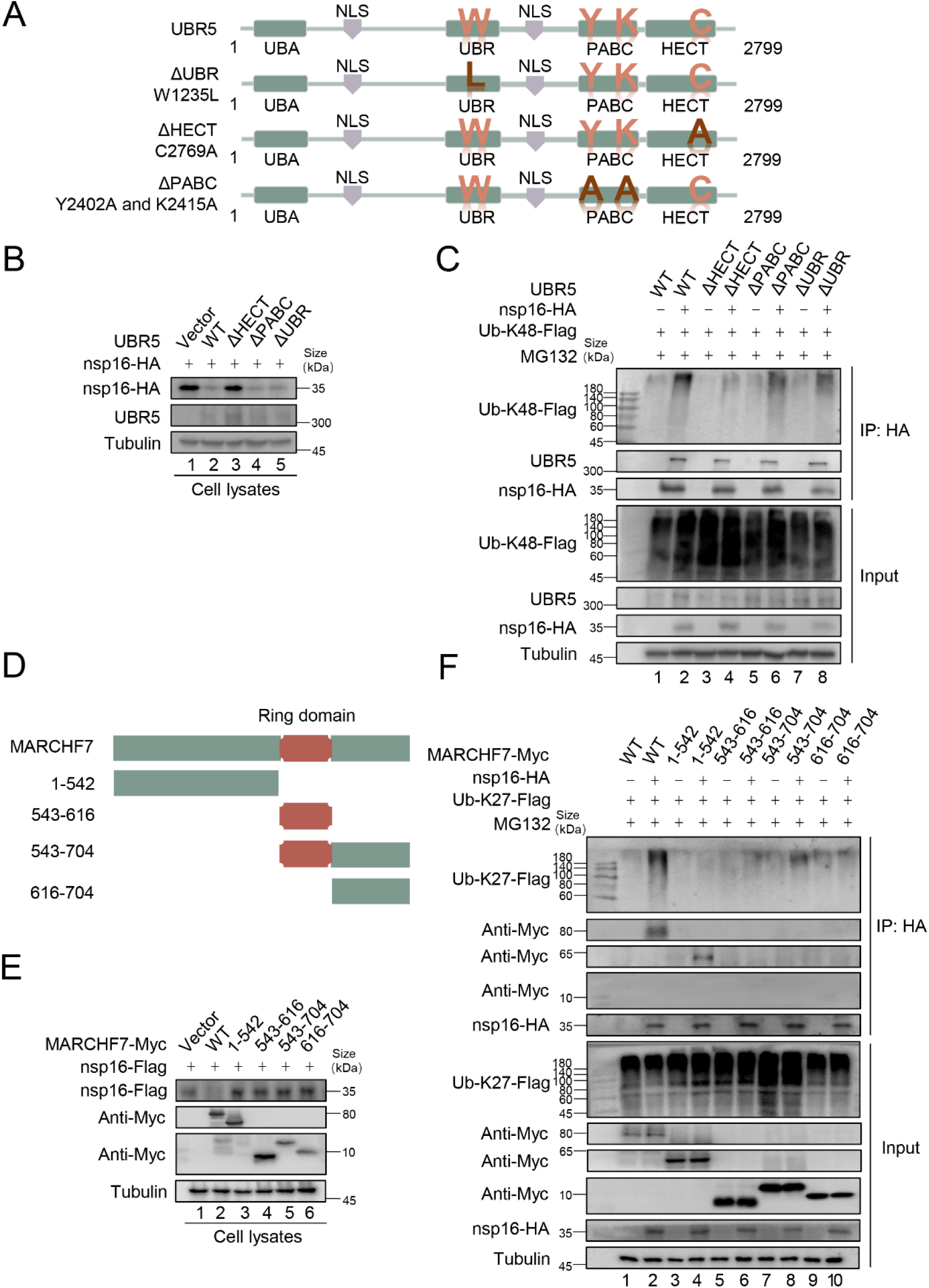
Domains in which MARCHF7 or UBR5 functions. A. The schematic represents UBR5 WT or mutants used in the study. B. The HECT domain of UBR5 is required for nsp16 degradation. After co-transfection with UBR5 WT or mutants and nsp16-HA, cells were harvested 48 hours later, and cell lysates were examined by IB. C. The HECT domain of UBR5 affects K48-type ubiquitin chain of nsp16.HEK293T cells were transfected with the assigned plasmids. After 36 hours, cells were treated with 10 µM MG132 for 12 hours, harvested, and cell lysates were incubated with protein G agarose beads conjugated with anti-HA antibodies. Cell lysates and precipitated samples were analyzed by IB. D. The schematic represents wild-type and truncated forms of MARCHF7 used in the study. E. Only MARCHF7 wild-type degraded nsp16. F. The N-terminal region of MARCHF7 interacted with nsp16, and only the wild type could catalyze the K27-type ubiquitin chain of nsp16. Figure 4—figure supplement 2-source data 1. PDF file containing original western blots for Figure 4—figure supplement 2B, C, E, and F, indicating the relevant bands and treatments. Figure 4 — figure supplement 2-source data 2. Original files for western blot analysis displayed in Figure 4—figure supplement 2B, C, E, and F.

**Figure 5—figure supplement 1.**
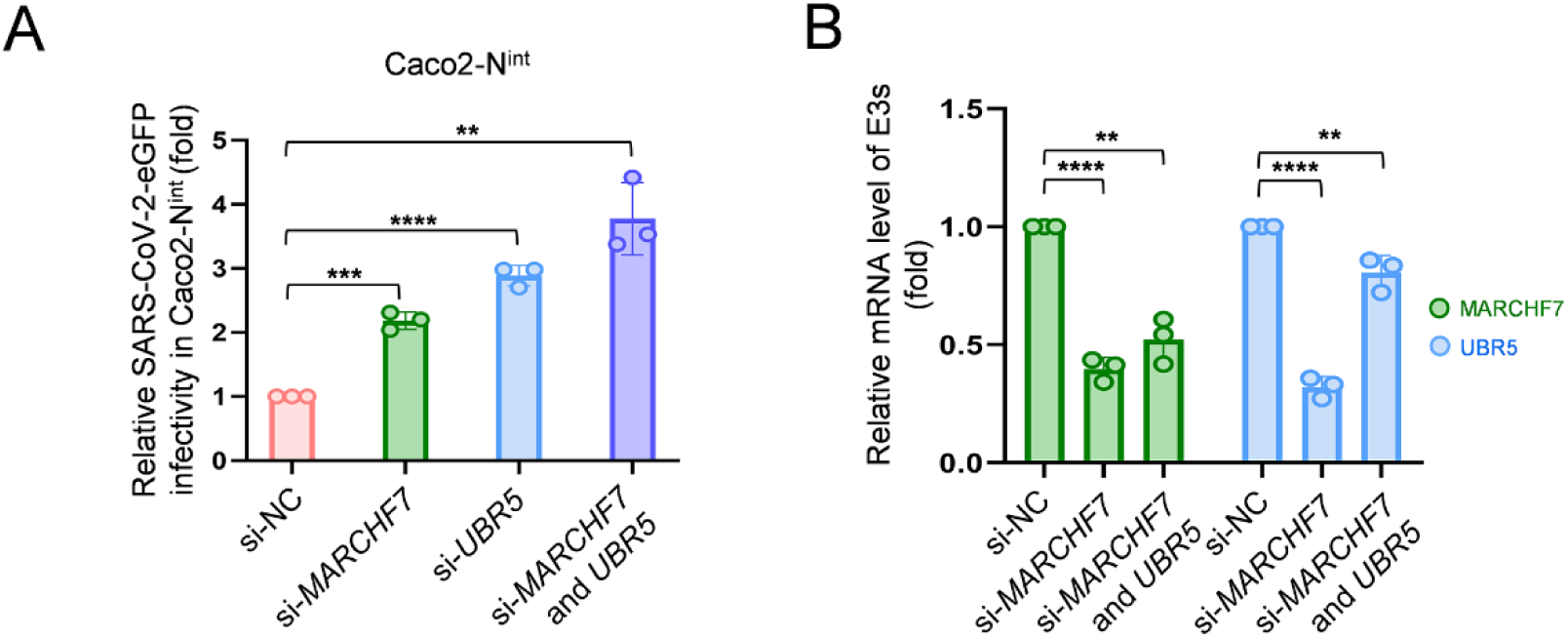
Effect of MARCHF7 or UBR5 on SARS-CoV-2 trVLP. (A-B). Knocking down *MARCHF7* or *UBR5* enhances SARS-CoV-2 trVLP infectivity. *MARCHF7* or *UBR5* was knocked down by siRNA in Caco2 cells with stable expression of SARS-CoV-2 N protein. Twenty-four hours later, cells were infected with SARS-CoV-2 virus-like-particles (MOI:0.1), the medium was changed two hours after infection, and the eGFP-positive cells were detected by flow cytometry 48 hours later (A). Protein content was determined by RT-qPCR (B). Figure 5—figure supplement 1-source data 1. Numerical data obtained during experiments represented in Figure 5—figure supplement 1.

**Figure 6—figure supplement 1.**
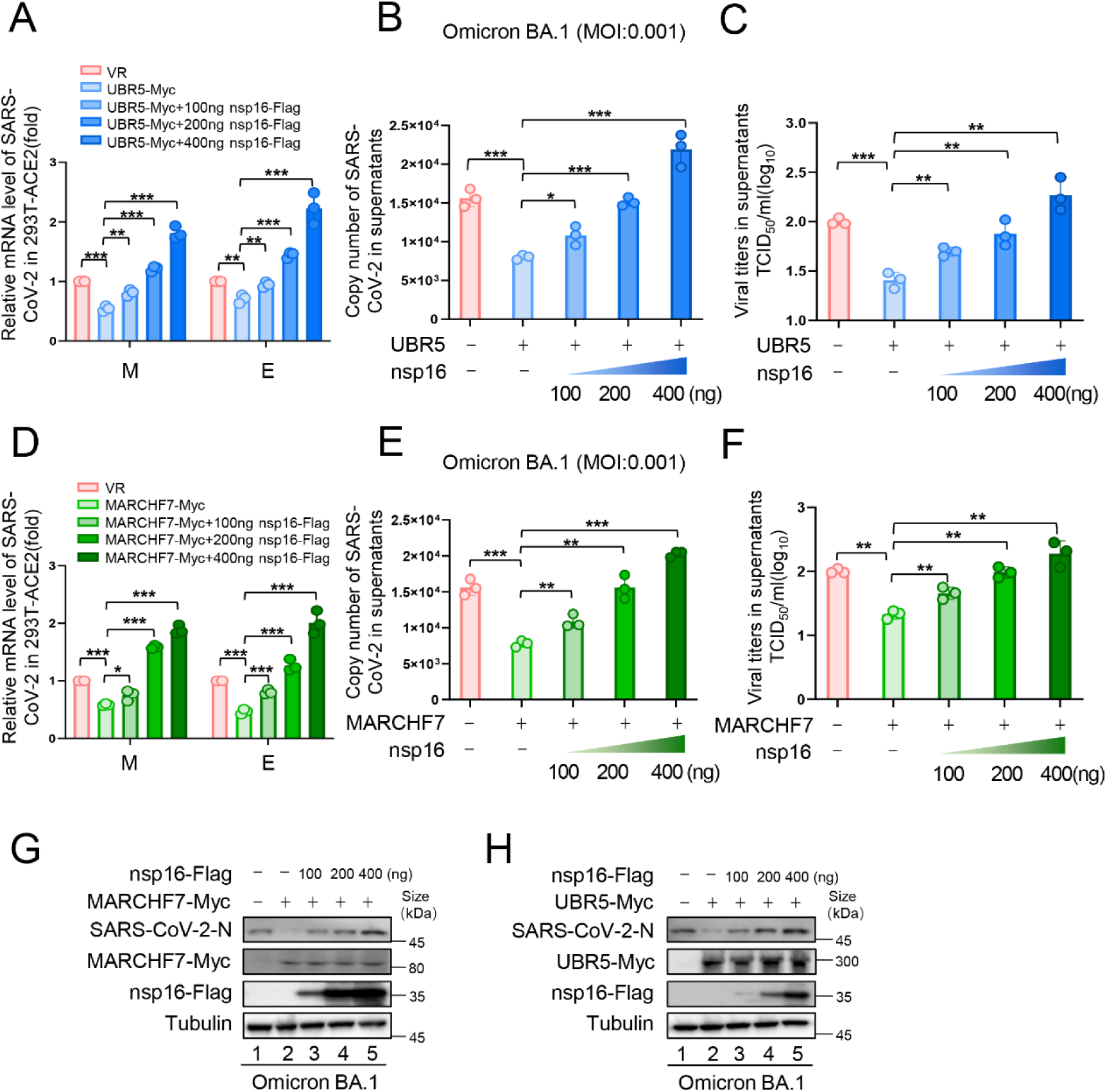
Effect of MARCHF7 or UBR5 on Omicron BA.1 infectivity. (A-H). Showing data related to infection with Omicron BA.1. The experimental procedure was the same as Figure 6. Data are representative of three independent experiments and shown as average ± SD (n = 3). Significance was determined by one-way ANOVA, followed by a Tukey multiple comparisons posttest. P > 0.05; **P < 0.01; ***P < 0.001. Figure 6—figure supplement 1-source data 1. PDF file containing original western blots for Figure 6—figure supplement 1G and H, indicating the relevant bands and treatments. Figure 6 — figure supplement 1-source data 2. Original files for western blot analysis displayed in Figure 6—figure supplement 1G and H. Figure 6—figure supplement 1-source data 3. Numerical data obtained during experiments represented in Figure 6—figure supplement 1.

**Figure 6—figure supplement 2.**
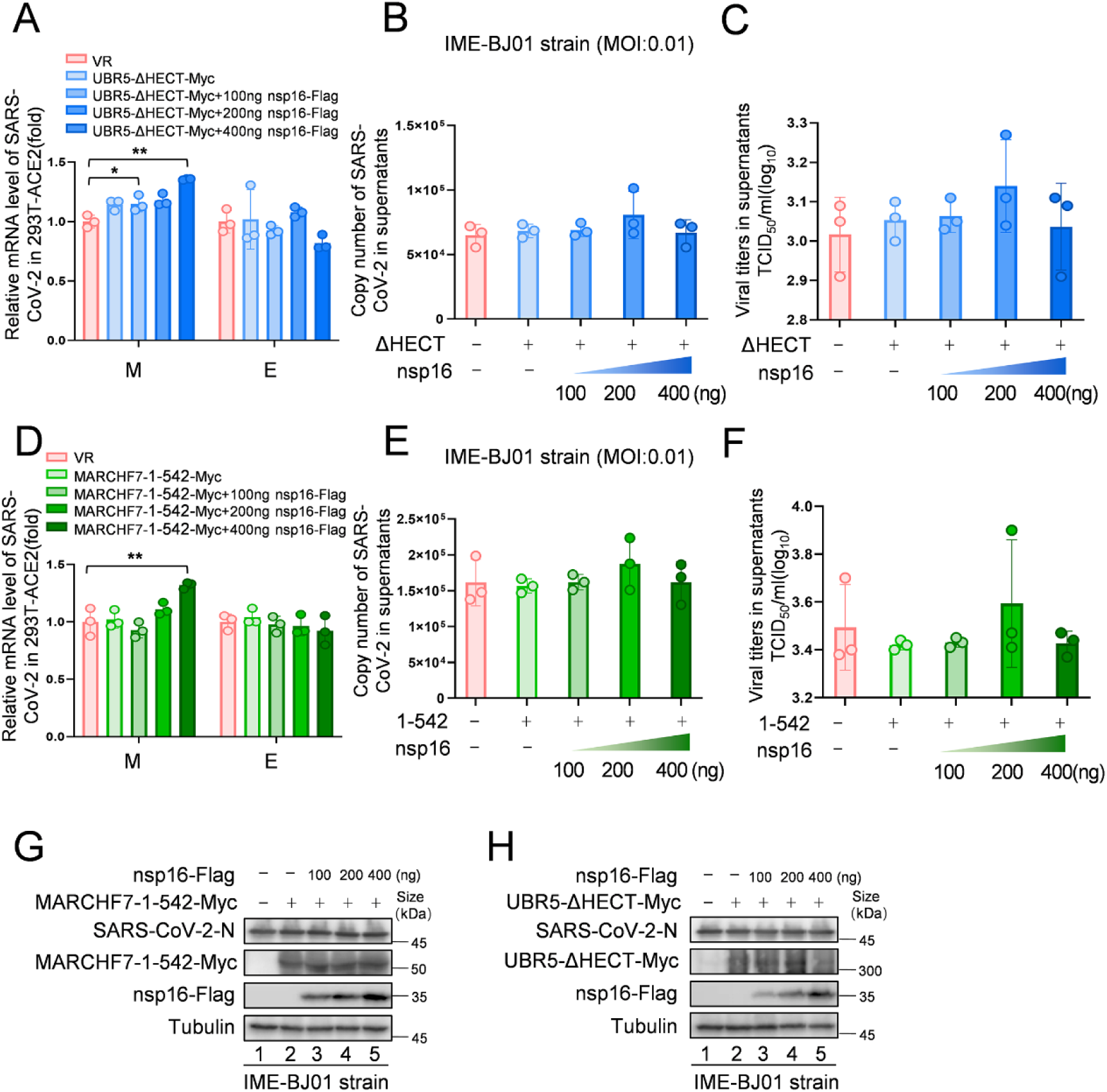
The enzyme activity-deficient mutants do not exhibit antiviral activity, and overexpression of nsp16 does not promote viral replication. (A-H) In 293T-ACE2 cells, the RING domain deletion mutant of MARCHF7 (MARCHF7-aa 1-542) or the HECT domain inactivated mutant of UBR5 (UBR5-ΔHECT) were transfected, along with a gradient of nsp16-Flag overexpression. The cells were infected with the IME-BJ01 strain (MOI: 0.01), medium was changed 2 hours post-infection, and cells and supernatants were collected 48 hours after infection. Figure 6—figure supplement 2-source data 1. PDF file containing original western blots for Figure 6—figure supplement 2G and H, indicating the relevant bands and treatments. Figure 6 — figure supplement 2-source data 2. Original files for western blot analysis displayed in Figure 6—figure supplement 2G and H. Figure 6—figure supplement 2-source data 3. Numerical data obtained during experiments represented in Figure 6—figure supplement 2.

**Figure 6—figure supplement 3.**
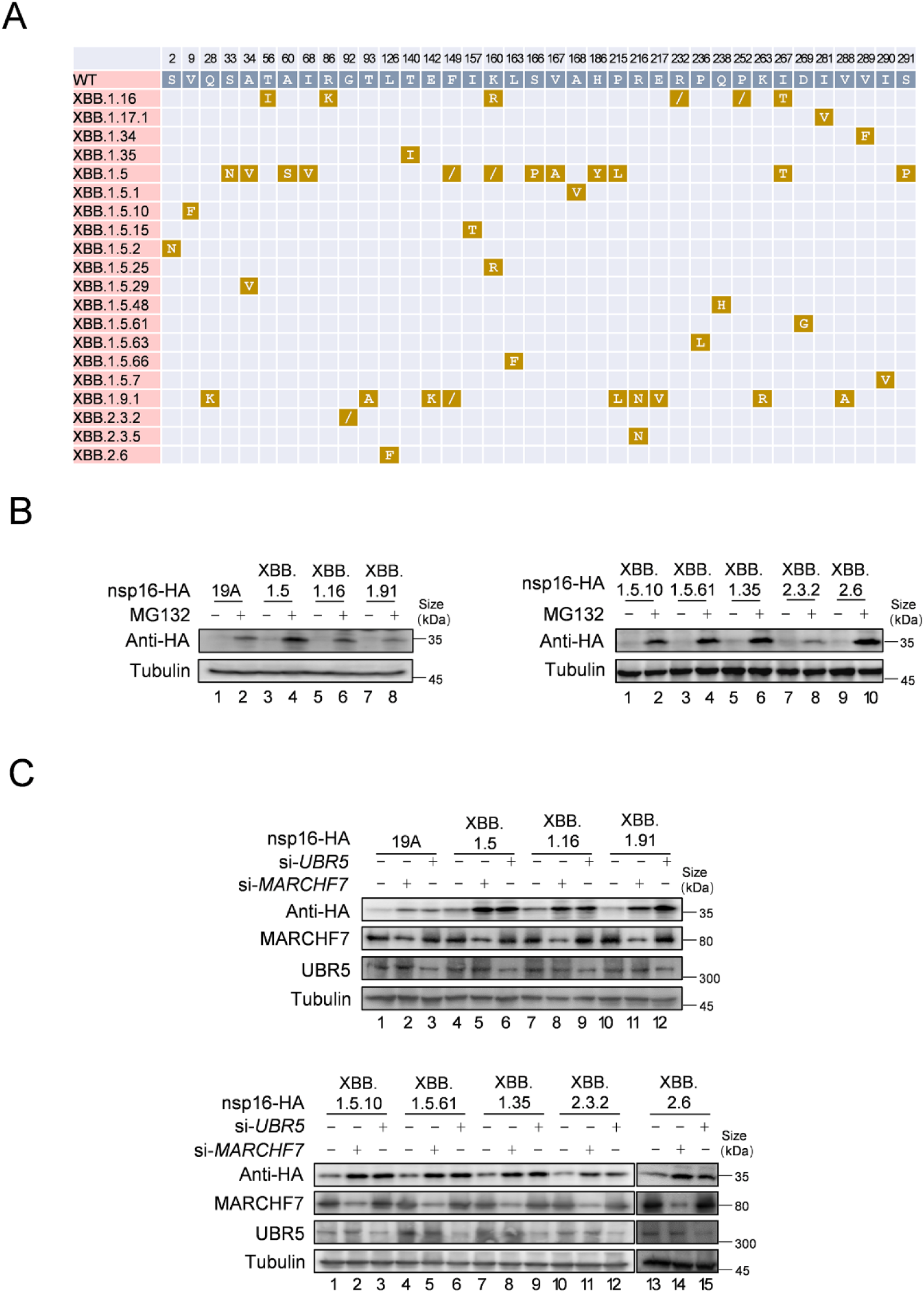
MARCHF7 or UBR5 have effects on mutant of nsp16 in different subtypes of SARS-CoV-2. A. This diagram shows the mutation of nsp16 in different virus subtypes. The amino acids sequences of different SARS-CoV-2 strains were obtained from NCBI, and the amino acids sequences of nsp16 of different strains were compared by DNAMAN software. B. Nsp16 mutants can still be regulated by MG132. The mutated nsp16 plasmids were transfected into HEK293T cells. After 36 hours of culture, cells were treated with 10 µm MG132 or DMSO, harvested 12 hours later, and cell lysates were examined by IB. C. MARCHF7 or UBR5 can degrade nsp16 mutants. After transfecting *MARCHF7* or *UBR5* siRNA and the mutated nsp16 plasmids, the cells were harvested 48 h later. The cell lysates were detected by IB. Figure 6—figure supplement 3-source data 1. PDF file containing original western blots for Figure 6—figure supplement 3B and C, indicating the relevant bands and treatments. Figure 6 — figure supplement 3-source data 2. Original files for western blot analysis displayed in Figure 6—figure supplement 3B and C.

**Figure 6—figure supplement 4.**
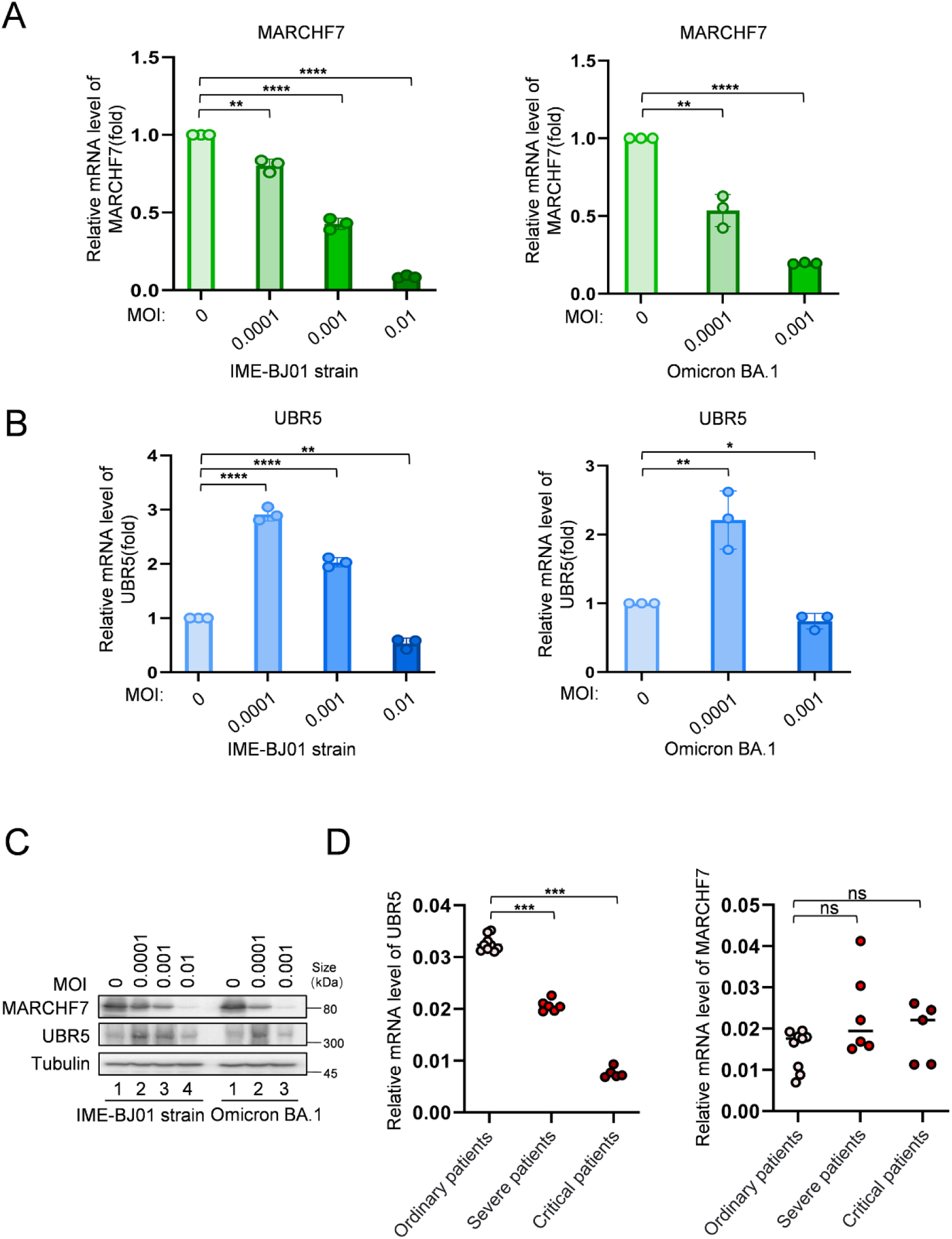
The expression level of MARCHF7 was negatively correlated with the viral titer, while the expression level of UBR5 was increased at low titer and decreased at high titer. (A-C). The protein and mRNA levels of MARCHF7 or UBR5 upon infection with different titers. Endogenous MARCHF7 and UBR5 RNA levels were detected by RT-qPCR 48 hours after infection with different titers of IME-BJ01 strain (MOI:0, 0.0001, 0.001, 0.01) or omicron BA.1 strain (MOI:0, 0.0001, 0.001). Protein levels were examined by IB. D. The expression level of UBR5 was negatively correlated with the severity of the disease but MARCHF7 expression levels were not. PBMC cells were extracted from common, severe and critical COVID-19 patients. RT-qPCR was used to detect the mRNA level of UBR5 or MARCHF7 in patients. Significance was determined by one-way ANOVA, followed by a Tukey multiple comparisons posttest. ns, P > 0.05; **P < 0.01; ***P < 0.001. Figure 6—figure supplement 4-source data 1. PDF file containing original western blots for Figure 6—figure supplement 4C, indicating the relevant bands and treatments. Figure 6 — figure supplement 4-source data 2. Original files for western blot analysis displayed in Figure 6—figure supplement 4C. Figure 6—figure supplement 4-source data 3. Numerical data obtained during experiments represented in Figure 6—figure supplement 4.

**Figure 8—figure supplement 1.**
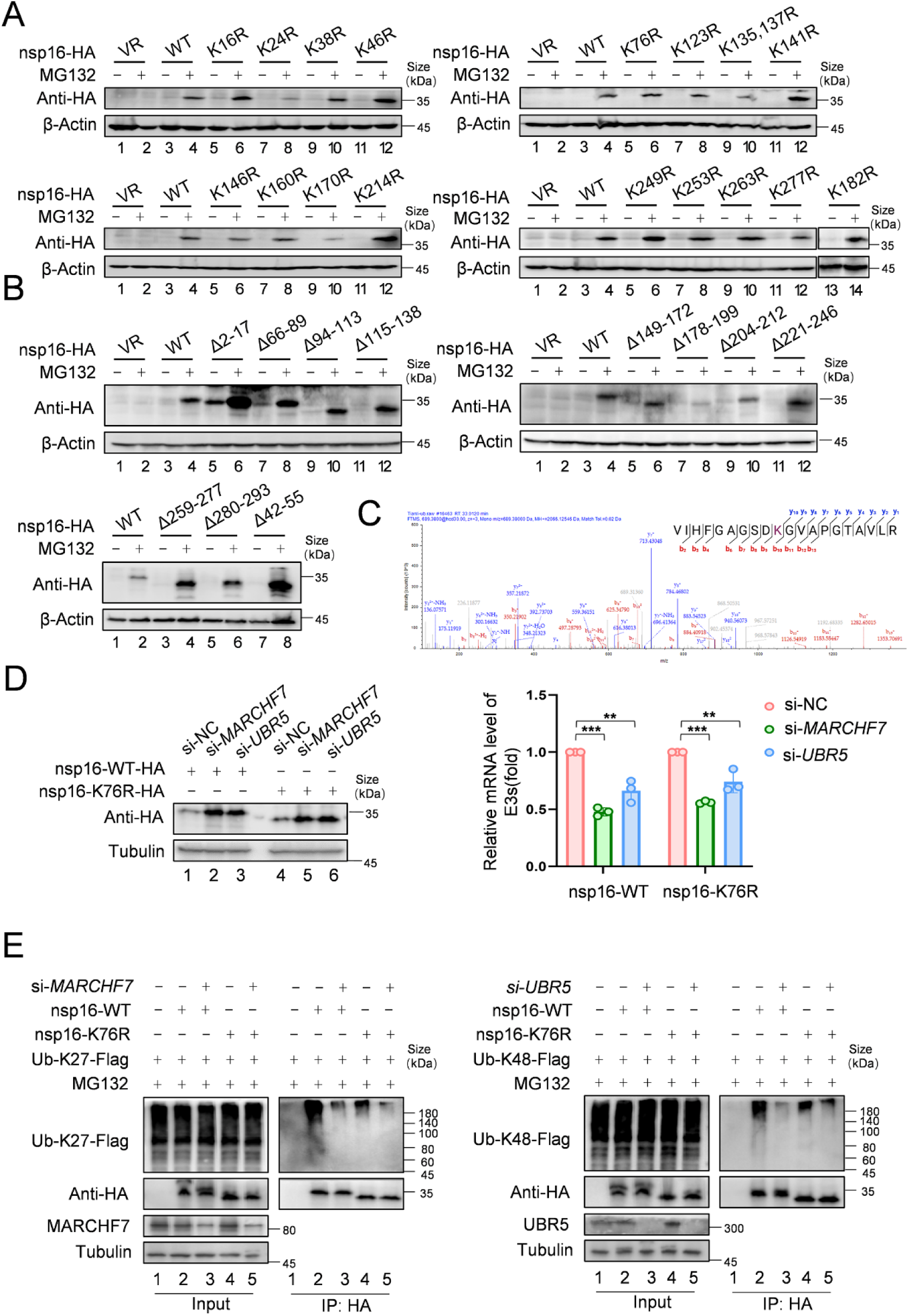
Identification of the ubiquitination modification site of nsp16 protein. A. Single lysine mutation of nsp16 protein can be restored by MG132. Single lysine mutants of nsp16 protein were obtained through mutagenesis and overexpressed in 293T cells. After 36 hours, cells were treated with either MG132 or DMSO for 16 hours and then harvested. Protein levels were detected by IB using the HA antibody. B. nsp16 protein truncates can be restored by MG132. nsp16 protein truncates were obtained through structural analysis and mutations. IB was performed to determine whether the truncates could be restored by MG132. C. The mass spectrometry analysis identified the ubiquitination modification site at lysine 76. nsp16-Flag was overexpressed in 293T cells, followed by MG132 treatment and cell harvest. nsp16 protein was enriched using Flag antibody-conjugated protein-G beads. Flag-peptide competition was used to obtain the nsp16-containing solution. The protein and ubiquitination status were visualized by SDS-PAGE and Coomassie staining. Mass spectrometry was used for further analysis. D. Degradation of nsp16-K76R is still regulated by MARCHF7 or UBR5. A plasmid with a mutation at lysine 76 of nsp16 to arginine (nsp16-K76R) was constructed. MARCHF7 or UBR5 was knocked down by siRNA in 293T cells. nsp16-WT or nsp16-K76R was transfected the next day, and cells were harvested 48 hours later. Protein levels were detected by IB. E. Ubiquitination levels of nsp16-K76R are reduced but still regulated by MARCHF7 or UBR5. MARCHF7 or UBR5 was knocked down using siRNA in 293T cells. The cells were co-transfected with Ub-K27 or K48, and nsp16-WT or nsp16-K76R mutant. Cells were harvested 48 hours later, with MG132 treatment 16 hours before harvesting. Co-IP experiments were performed to analyze the ubiquitination status of nsp16-WT or its mutant. Figure 8—figure supplement 1-source data 1. PDF file containing original western blots for Figure 8—figure supplement 1A, B, D, and E, indicating the relevant bands and treatments. Figure 8—figure supplement 1-source data 2. Original files for western blot analysis displayed in Figure 8—figure supplement 1A, B, D, and E. Figure 8—figure supplement 1-source data 3. Numerical data obtained during experiments represented in Figure 8—figure supplement 1.

## Additional files

**Supplementary file 1: Primers used in this study**

**Supplementary file 2: Patients clinical information in this study**

Dear Editor,

Thank you very much for your valuable feedback and for pointing out the formatting inconsistencies in our manuscript. We sincerely appreciate the time and effort you and the reviewers have devoted to improving the quality of our submission.

We have addressed the requested revisions as follows:

1. Thank you for providing source data for the gels / blots. Please could you reorganise this as follows: For each individual figure and individual figure supplement please provide two zipped folders, the first containing (1) the original files of the full raw uncropped, unedited gels or blots and; the second containing (2) figures with the uncropped gels or blots with the relevant bands clearlylabelled. These should be clearly labelled to indicate which figure/figure supplement each zipped folder relates to (Figure 1-source data 1, Figure 1-source data 2, Figure 3-source data 1, Figure 3-source data 1, Figure 3-figure supplement 1-source data 1, Figure 3-figure supplement 1-source data2, etc.) and should be accompanied by descriptive titles in the article file. Response: We have now reorganized the original gel/blot data as requested. For each main figure and figure supplement, we have provided two separate zipped folders: One containing the original, uncropped, unedited gel or blot images in their native formats. Another with uncropped images labeled to indicate the relevant bands used in the figures. Each folder is clearly named to correspond with the respective figure or figure supplement (e.g., Figure 1-source data 1, Figure 1-source data 2, Figure 3-figure supplement 1-source data 1, etc.), and appropriate descriptive titles have been included in the article file accordingly.
2. We notice you have included an appendix which consists of only figures and/or tables. As there is no associated appendix text, we would like to ask you to remove the appendix section and modify how these figures/tables are included: For Figures, please move these to the main article, either as additional main figures or as figure supplements to main figures. Please ensure each figure/figure supplement is uploaded individually using the relevant file type and the labelling in the submission form and any citations in the articlefile and response to reviews are in the format “Figure 1”, “Figure 2”, “Figure 3” etc. for main figures and “Figure 1-figure supplement 1”, “Figure 1-figure supplement 2”, “Figure 2-figure supplement 1”. For the tables: if these should be included as main display items, please instead embed these in the main article file in an editable format, updating the label and any references in the article file and response to reviews to follow the format “Table 1”, “Table 2” etc. If the tables are notappropriate to be displayed in the main text, could these instead be uploaded as supplementary files or source data files, as appropriate, and in a searchable/editable format (e.g. as excel files)? Any supplementary files should be labelled in the submission form and cited in the article file andresponse to reviews in the format “Supplementary file 1”, “Supplementary file 2” etc. Source data files should be linked to a main figure or table or figure supplement where possible e.g. Figure 1-source data 1, Figure 1-source data 2, Table 1-source data 1, Figure 1-figure supplement 1-source data 1. Response: We have removed the Appendix section as it only contained figures and tables without any associated text. The figures from the Appendix have been relocated as follows: Appendix Figure 1 has been incorporated into Figure 2-figure supplement 1C. Appendix Figure 2 has been incorporated into Figure 8-figure supplement 1. We have ensured that all figure legends are now properly placed and labeled. Furthermore, the tables previously included in the Appendix have been uploaded as supplementary files in an editable format, and they are now appropriately cited in the article text as Supplementary file 1, and Supplementary file 2. Thank you again for your guidance and support throughout this process.

## Notes

Competing Interests: The authors declare no conflicts of interest.

### Competing Interest Statement

The authors have declared no competing interest.

### Summary of Updates

We have removed the Appendix section as it only contained figures and tables without any associated text. The figures from the Appendix have been relocated as follows: Appendix Figure 1 has been incorporated into Figure 2-figure supplement 1C. Appendix Figure 2 has been incorporated into Figure 8-figure supplement 1. We have ensured that all figure legends are now properly placed and labeled. Furthermore, the tables previously included in the Appendix have been uploaded as supplementary files in an editable format, and they are now appropriately cited in the article text as Supplementary file 1, and Supplementary file 2.

## References

Balieiro, A. M., Anunciação, E. L., Costa, C. H., Qayed, W. S., & Silva, J. R. A. (2022). Computational Analysis of SAM Analogs as Methyltransferase Inhibitors of nsp16/nsp10 Complex from SARS-CoV-2. International Journal of Molecular Sciences, 23(22), 13972.

Banerjee, A. K., Blanco, M. R., Bruce, E. A., Honson, D. D., Chen, L. M., Chow, A., Bhat, P., Ollikainen, N., Quinodoz, S. A., & Loney, C. (2020). SARS-CoV-2 disrupts splicing, translation, and protein trafficking to suppress host defenses. Cell, 183(5), 1325–1339. e1321.

Bonamassa, B., Hai, L., & Liu, D. (2011). Hydrodynamic gene delivery and its applications in pharmaceutical research. Pharm Res, 28(4), 694–701. 10.1007/s11095-010-0338-9

Cai, B., Zhao, J., Zhang, Y., Liu, Y., Ma, C., Yi, F., Zheng, Y., Zhang, L., Chen, T., & Liu, H. (2022). USP5 attenuates NLRP3 inflammasome activation by promoting autophagic degradation of NLRP3. Autophagy, 18(5), 990–1004.

Chen, X., Tian, L., Zhang, L., Gao, W., Yu, M., Li, Z., & Zhang, W. (2024). Deubiquitinase USP39 promotes SARS-CoV-2 replication by deubiquitinating and stabilizing the envelope protein. Antiviral Res, 221, 105790. 10.1016/j.antiviral.2023.105790

Decroly, E., Debarnot, C., Ferron, F., Bouvet, M., Coutard, B., Imbert, I., Gluais, L., Papageorgiou, N., Sharff, A., & Bricogne, G. (2011). Crystal structure and functional analysis of the SARS-coronavirus RNA cap 2′-O-methyltransferase nsp10/nsp16 complex. PLoS pathogens, 7(5), e1002059.

Fan, L., Zhou, Y., Wei, X., Feng, W., Guo, H., Li, Y., Gao, X., Zhou, J., Wen, Y., Wu, Y., Shen, X., Liu, L., Xu, G., & Zhang, Z. (2024). The E3 ligase TRIM22 restricts SARS-CoV-2 replication by promoting proteasomal degradation of NSP8. mBio, 15(2), e0232023. 10.1128/mbio.02320-23

Gao, W., Wang, L., Cui, W., Wang, H., Huang, G., Li, Z., Li, G., & Zhang, W. (2024). Deubiquitinase USP1 regulates sarbecovirus ORF6 protein function. J Virol, 98(1), e0143723. 10.1128/jvi.01437-23

Gao, W., Wang, L., Ju, X., Zhao, S., Li, Z., Su, M., Xu, J., Wang, P., Ding, Q., & Lv, G. (2022). The deubiquitinase USP29 promotes SARS-CoV-2 virulence by preventing proteasome degradation of ORF9b. Mbio, 13(3), e01300–01322.

Grice, G. L., & Nathan, J. A. (2016). The recognition of ubiquitinated proteins by the proteasome. Cellular and molecular life sciences, 73, 3497–3506.

Guo, G., Gao, M., Gao, X., Zhu, B., Huang, J., Luo, K., Zhang, Y., Sun, J., Deng, M., & Lou, Z. (2021). SARS-CoV-2 non-structural protein 13 (nsp13) hijacks host deubiquitinase USP13 and counteracts host antiviral immune response. Signal transduction and targeted therapy, 6(1), 119.

Hojyo, S., Uchida, M., Tanaka, K., Hasebe, R., Tanaka, Y., Murakami, M., & Hirano, T. (2020). How COVID-19 induces cytokine storm with high mortality. Inflamm Regen, 40, 37. 10.1186/s41232-020-00146-3

Hu, B., Huang, S., & Yin, L. (2021). The cytokine storm and COVID-19. J Med Virol, 93(1), 250–256. 10.1002/jmv.26232

Ju, X., Zhu, Y., Wang, Y., Li, J., Zhang, J., Gong, M., Ren, W., Li, S., Zhong, J., Zhang, L., Zhang, Q. C., Zhang, R., & Ding, Q. (2021). A novel cell culture system modeling the SARS-CoV-2 life cycle. PLoS Pathog, 17(3), e1009439. 10.1371/journal.ppat.1009439

Kamimura, K., Kanefuji, T., Yokoo, T., Abe, H., Suda, T., Kobayashi, Y., Zhang, G., Aoyagi, Y., & Liu, D. (2014). Safety assessment of liver-targeted hydrodynamic gene delivery in dogs. PLoS One, 9(9), e107203. 10.1371/journal.pone.0107203

Kim, J. G., Shin, H.-C., Seo, T., Nawale, L., Han, G., Kim, B. Y., Kim, S. J., & Cha-Molstad, H. (2021). Signaling pathways regulated by UBR box-containing E3 ligases. International journal of molecular sciences, 22(15), 8323.

Klima, M., Khalili Yazdi, A., Li, F., Chau, I., Hajian, T., Bolotokova, A., Kaniskan, H. Ü., Han, Y., Wang, K., & Li, D. (2022). Crystal structure of SARS-CoV-2 nsp10–nsp16 in complex with small molecule inhibitors, SS148 and WZ16. Protein Science, 31(9), e4395.

Li, Z., Hao, P., Zhao, Z., Gao, W., Huan, C., Li, L., Chen, X., Wang, H., Jin, N., & Luo, Z.-Q. (2023). The E3 ligase RNF5 restricts SARS-CoV-2 replication by targeting its envelope protein for degradation. Signal Transduction and Targeted Therapy, 8(1), 53.

Liang, X., Xiao, J., Li, X., Liu, Y., Lu, Y., Wen, Y., Li, Z., Che, X., Ma, Y., & Zhang, X. (2022). A C-terminal glutamine recognition mechanism revealed by E3 ligase TRIM7 structures. Nature Chemical Biology, 18(11), 1214–1223.

Lin, S., Chen, H., Ye, F., Chen, Z., Yang, F., Zheng, Y., Cao, Y., Qiao, J., Yang, S., & Lu, G. (2020). Crystal structure of SARS-CoV-2 nsp10/nsp16 2′-O-methylase and its implication on antiviral drug design. Signal transduction and targeted therapy, 5(1), 131.

Lugari, A., Betzi, S., Decroly, E., Bonnaud, E., Hermant, A., Guillemot, J.-C., Debarnot, C., Borg, J.-P., Bouvet, M., & Canard, B. (2010). Molecular mapping of the RNA Cap 2 ′ -O-methyltransferase activation interface between severe acute respiratory syndrome coronavirus nsp10 and nsp16. Journal of Biological Chemistry, 285(43), 33230–33241.

Mahajan, S., Sharma, G. K., Bora, K., & Pattnaik, B. (2021). Identification of novel interactions between host and non-structural protein 2C of foot-and-mouth disease virus. J Gen Virol, 102(3). 10.1099/jgv.0.001577

Maimaitiyiming, Y., Yang, T., Wang, Q. Q., Feng, Y., Chen, Z., Björklund, M., Wang, F., Hu, C., Hsu, C.-H., & Naranmandura, H. (2022). Heat treatment promotes ubiquitin-mediated proteolysis of SARS-CoV-2 RNA polymerase and decreases viral load. Research.

Makaremi, S., Asgarzadeh, A., Kianfar, H., Mohammadnia, A., Asghariazar, V., & Safarzadeh, E. (2022). The role of IL-1 family of cytokines and receptors in pathogenesis of COVID-19. Inflamm Res, 71(7-8), 923–947. 10.1007/s00011-022-01596-w

McClellan, A. J., Laugesen, S. H., & Ellgaard, L. (2019). Cellular functions and molecular mechanisms of non-lysine ubiquitination. Open Biology, 9(9), 190147.

Melo-Filho, C. C., Bobrowski, T., Martin, H.-J., Sessions, Z., Popov, K. I., Moorman, N. J., Baric, R. S., Muratov, E. N., & Tropsha, A. (2022). Conserved coronavirus proteins as targets of broad-spectrum antivirals. Antiviral Research, 204, 105360.

Muñoz-Escobar, J., Matta-Camacho, E., Kozlov, G., & Gehring, K. (2015). The MLLE domain of the ubiquitin ligase UBR5 binds to its catalytic domain to regulate substrate binding. Journal of Biological Chemistry, 290(37), 22841–22850.

Nathan, J. A., Sengupta, S., Wood, S. A., Admon, A., Markson, G., Sanderson, C., & Lehner, P. J. (2008). The ubiquitin E3 ligase MARCH7 is differentially regulated by the deubiquitylating enzymes USP7 and USP9X. Traffic, 9(7), 1130–1145.

Park, G. J., Osinski, A., Hernandez, G., Eitson, J. L., Majumdar, A., Tonelli, M., Henzler-Wildman, K., Pawłowski, K., Chen, Z., & Li, Y. (2022). The mechanism of RNA capping by SARS-CoV-2. Nature, 609(7928), 793–800.

Qiao, X., Liu, Y., Prada, M. L., Mohan, A. K., Gupta, A., Jaiswal, A., Sharma, M., Merisaari, J., Haikala, H. M., & Talvinen, K. (2020). UBR5 is coamplified with MYC in breast tumors and encodes an ubiquitin ligase that limits MYC-dependent apoptosis. Cancer research, 80(7), 1414–1427.

Raper, S. E., Chirmule, N., Lee, F. S., Wivel, N. A., Bagg, A., Gao, G. P., Wilson, J. M., & Batshaw, M. L. (2003). Fatal systemic inflammatory response syndrome in a ornithine transcarbamylase deficient patient following adenoviral gene transfer. Mol Genet Metab, 80(1-2), 148–158. 10.1016/j.ymgme.2003.08.016

Rosas-Lemus, M., Minasov, G., Shuvalova, L., Inniss, N. L., Kiryukhina, O., Brunzelle, J., & Satchell, K. J. (2020). High-resolution structures of the SARS-CoV-2 2′-O-methyltransferase reveal strategies for structure-based inhibitor design. Science signaling, 13(651), eabe1202.

Russ, A., Wittmann, S., Tsukamoto, Y., Herrmann, A., Deutschmann, J., Lagisquet, J., Ensser, A., Kato, H., & Gramberg, T. (2022). Nsp16 shields SARS–CoV-2 from efficient MDA5 sensing and IFIT1-mediated restriction. EMBO reports, 23(12), e55648.

Shearer, R. F., Frikstad, K.-A. M., McKenna, J., McCloy, R. A., Deng, N., Burgess, A., Stokke, T., Patzke, S., & Saunders, D. N. (2018). The E3 ubiquitin ligase UBR5 regulates centriolar satellite stability and primary cilia. Molecular biology of the cell, 29(13), 1542–1554.

Song, P., Li, W., Xie, J., Hou, Y., & You, C. (2020). Cytokine storm induced by SARS-CoV-2. Clin Chim Acta, 509, 280–287. 10.1016/j.cca.2020.06.017

Suda, T., Yokoo, T., Kanefuji, T., Kamimura, K., Zhang, G., & Liu, D. (2023). Hydrodynamic Delivery: Characteristics, Applications, and Technological Advances. Pharmaceutics, 15(4). 10.3390/pharmaceutics15041111

Wu, B., Song, M., Dong, Q., Xiang, G., Li, J., Ma, X., & Wei, F. (2022). UBR5 promotes tumor immune evasion through enhancing IFN-γ-induced PDL1 transcription in triple negative breast cancer. Theranostics, 12(11), 5086.

Xiang, G., Wang, S., Chen, L., Song, M., Song, X., Wang, H., Zhou, P., Ma, X., & Yu, J. (2022). UBR5 targets tumor suppressor CDC73 proteolytically to promote aggressive breast cancer. Cell Death & Disease, 13(5), 451.

Xu, G., Wu, Y., Xiao, T., Qi, F., Fan, L., Zhang, S., Zhou, J., He, Y., Gao, X., & Zeng, H. (2022). Multiomics approach reveals the ubiquitination-specific processes hijacked by SARS-CoV-2. Signal Transduction and Targeted Therapy, 7(1), 312.

Zhang, H., Zheng, H., Zhu, J., Dong, Q., Wang, J., Fan, H., Chen, Y., Zhang, X., Han, X., & Li, Q. (2021). Ubiquitin-modified proteome of SARS-CoV-2-infected host cells reveals insights into virus–host interaction and pathogenesis. Journal of Proteome Research, 20(5), 2224–2239.

Zhang, J., Cruz-Cosme, R., Zhuang, M.-W., Liu, D., Liu, Y., Teng, S., Wang, P.-H., & Tang, Q. (2020). A systemic and molecular study of subcellular localization of SARS-CoV-2 proteins. Signal transduction and targeted therapy, 5(1), 269.

Zhang, L., Hao, P., Chen, X., Lv, S., Gao, W., Li, C., Li, Z., & Zhang, W. (2024). CRL4B E3 ligase recruited by PRPF19 inhibits SARS-CoV-2 infection by targeting ORF6 for ubiquitin-dependent degradation. mBio, 15(2), e0307123. 10.1128/mbio.03071-23

Zhang, L., Wang, H., Tian, L., & Li, H. (2016). Expression of USP7 and MARCH7 is correlated with poor prognosis in epithelial ovarian cancer. The Tohoku journal of experimental medicine, 239(3), 165–175.

Zhang, X., Yang, Z., Pan, T., Sun, Q., Chen, Q., Wang, P. H., Li, X., & Kuang, E. (2023). SARS-CoV-2 Nsp8 suppresses MDA5 antiviral immune responses by impairing TRIM4-mediated K63-linked polyubiquitination. PLoS Pathog, 19(11), e1011792. 10.1371/journal.ppat.1011792

Zhao, K., Yang, Y., Zhang, G., Wang, C., Wang, D., Wu, M., & Mei, Y. (2018). Regulation of the Mdm2–p53 pathway by the ubiquitin E3 ligase MARCH 7. EMBO reports, 19(2), 305–319.

Zheng, C. (2021). The emerging roles of the MARCH ligases in antiviral innate immunity. International Journal of Biological Macromolecules, 171, 423–427.

Zhou, Y., Zheng, R., Liu, D., Liu, S., Disoma, C., Li, S., Liao, Y., Chen, Z., Du, A., & Dong, Z. (2022). UBR5 Acts as an Antiviral Host Factor against MERS-CoV via Promoting Ubiquitination and Degradation of ORF4b. Journal of Virology, 96(17), e00741–00722.

Züst, R., Cervantes-Barragan, L., Habjan, M., Maier, R., Neuman, B. W., Ziebuhr, J., Szretter, K. J., Baker, S. C., Barchet, W., & Diamond, M. S. (2011). Ribose 2′-O-methylation provides a molecular signature for the distinction of self and non-self mRNA dependent on the RNA sensor Mda5. Nature immunology, 12(2), 137–143.

